# An AI-guided signature reveals the nature of the shared proximal pathways of host immune response in MIS-C and Kawasaki disease

**DOI:** 10.1101/2021.04.11.439347

**Authors:** Debashis Sahoo, Gajanan D. Katkar, Chisato Shimizu, Jihoon Kim, Soni Khandelwal, Adriana H. Tremoulet, John Kanegaye, Pediatric Emergency Medicine Kawasaki Disease Research Group, Joseph Bocchini, Soumita Das, Jane C. Burns, Pradipta Ghosh

**Author notes:** Equal contribution (Debashis Sahoo, Gajanan D. Katkar, Chisato Shimizu). Consortia: Pediatric Emergency Medicine Kawasaki Disease Research Group (see Supplemental Online Material). **Correspondence to**: **Debashis Sahoo, Ph.D;** Associate Professor, Department of Pediatrics, University of California San Diego; 9500 Gilman Drive, MC 0703, Leichtag Building 132; La Jolla, CA 92093-0831., **Jane C. Burns, M.D.;** Director, Kawasaki Disease Research Center, Department of Pediatrics, University of California San Diego; 9500 Gilman Dr. MC 0641, La Jolla, CA 92093-0641, **Pradipta Ghosh, M.D.;** Professor, Departments of Medicine, and Cell and Molecular Medicine, University of California San Diego; 9500 Gilman Drive (MC 0651), George E. Palade Bldg, Rm 232, 239; La Jolla, CA 92093.

## Abstract

A significant surge in cases of multisystem inflammatory syndrome in children (MIS-C, also called Pediatric Inflammatory Multisystem Syndrome - PIMS) has been observed amidst the COVID-19 pandemic. MIS-C shares many clinical features with Kawasaki disease (KD), although clinical course and outcomes are divergent. We analyzed whole blood RNA sequences, serum cytokines, and formalin fixed heart tissues from these patients using a computational toolbox of two gene signatures, i.e., the 166-gene viral pandemic (ViP) signature, and its 20-gene severe (s)ViP subset that were developed in the context of SARS-CoV-2 infection and a 13-transcript signature previously demonstrated to be diagnostic for KD. Our analyses revealed that KD and MIS-C are on the same continuum of the host immune response as COVID-19. While both the pediatric syndromes converge upon an *IL15/IL15RA*-centric cytokine storm, suggestive of shared proximal pathways of immunopathogenesis, they diverge in other laboratory parameters and cardiac phenotypes. The ViP signatures also revealed unique targetable cytokine pathways in MIS-C, place MIS-C farther along in the spectrum in severity compared to KD and pinpoint key clinical (reduced cardiac function) and laboratory (thrombocytopenia and eosinopenia) parameters that can be useful to monitor severity.

## INTRODUCTION

Multisystem inflammatory syndrome in children^1^ (MIS-C; initially named Pediatric Inflammatory Multisystem Syndrome Temporally associated with SARS-CoV-2, PIMS-TS)^2^ is a rare but severe condition that occurs in children and adolescents ~4–6 weeks after exposure to SARS-CoV-2. First reported in April 2020 in a cluster of children in the United Kingdom^3^, followed by other regions of the world ^4^, the syndrome is characterized by fever, and variously accompanied by rash, conjunctival injection, gastrointestinal symptoms, shock, and elevated markers of inflammation and antibodies to SARS-CoV-2 in the majority of patients. Myocardial dysfunction and coronary arterial dilation may resemble those seen in another uncommon childhood condition, Kawasaki Disease (KD). KD is an acute inflammatory disorder predominantly seen in young children. Since it was first described in Japan^5^ in 1967, KD has emerged as the most common cause of pediatric acquired heart disease in the developed world^6^. Little is known about the definitive triggers of KD; what is most widely accepted is that KD is largely an immune response to a plethora of infectious or environmental stimuli including viruses, fungi (e.g., *Candida* sp.), and bacteria ^7–9^. The host genetic background appears to shape this idiosyncratic inflammatory response to an environmental antigen exposure^9^.

On May 14, 2020, the CDC published an online Health Advisory that christened this condition as Multisystem Inflammatory Syndrome in Children (MIS-C) and outlined a case definition^10^. Since then, as the COVID-19 pandemic spread across many countries, cases of MIS-C soared, with features of shock and cardiac involvement requiring ionotropic support [in the critical care setting]. But distinguishing MIS-C from KD, KD shock syndrome^11^, and other severe infectious or inflammatory conditions remains a challenge. The need for early diagnostic and prognostic markers of disease severity remains unmet; such markers could objectively guide decisions regarding the appropriateness of the level of care and the timing of initiation of life-saving supportive and therapeutic measures.

As for the immunopathogenesis of MIS-C, limited but key insights have emerged rapidly, most of which focus on the differences between MIS-C and KD. For example, Gruber et al.,^12^ and Consiglio et al.,^13^ showed that the inflammatory response in MIS-C shares several features with KD, but also differs from this condition with respect to T cell subsets^13^. These conclusions were generally supported by two other studies, by Vella et al.,^14^ and Ramaswamy et al.^15^ who also showed that severe MIS-C patients displayed skewed memory T cell TCR repertoires and autoimmunity characterized by endothelium-reactive IgG. Finally, Carter et al.,^16^ reported activation of CD4^+^CCR7^+^ T cells and γδ T cell subsets in MIS-C, which had not been reported in KD, which made them conclude that MIS-C may be a distinct immunopathogenic illness. While these studies further our understanding of MIS-C and the major conclusions of these studies are comprehensively reviewed elsewhere^17^, it is noteworthy that each of these studies had some notable limitations— (i) in Gruber et al.,^12^ most of the MIS-C subjects were on immunomodulatory medications when samples were drawn; (ii) in Vella et al.,^14^ absence of contemporaneously analyzed healthy pediatric samples which were not available during the early phase of the pandemic; (iii) in Carter et al.,^16^ KD subjects were not concurrently studied and the authors themselves acknowledged that such side-by-side immunophenotyping of MIS-C and KD would be necessary to draw conclusions convincingly regarding similarities and differences between these two syndromes; (iv) absence of validation studies in independent cohorts in them all.

We recently showed that a 166-gene signature is conserved in all *viral p*andemics (ViP), including COVID-19, and a subset of 20-genes within that signature that classifies disease severity^18^. In the absence of a sufficiently large number of COVID-19 datasets at the onset of the COVID-19 pandemic, these ViP signatures were trained on datasets from the pandemics of the past (Influenza and avian flu) and used without further training to prospectively analyze the samples from the current pandemic (i.e., COVID-19). The ViP signatures appeared to capture the invariant host response shared between all viral pandemics, including COVID-19. Here we used the ViP signatures as a starting computational framework to navigate the syndrome of MIS-C that is still a relatively poorly understood entity, and to interrogate concurrently the shared and unique features in MIS-C and KD.

## RESULTS

### A gene signature seen in COVID-19 is also induced in KD, and tracks disease severity

We sought to define the host immune response in KD and compare that to COVID-19 using an artificial intelligence (AI)-based approach. To this end, we took advantage of a recently identified analysis of the host immune response in COVID-19 in which over 45,000 transcriptomic datasets of viral pandemics were analyzed to extract a 166-gene signature^18^ (summarized in **Figure 1A**). Because publicly available transcriptomic datasets from SARS-CoV-2-infected patients were still relatively few at the onset of the pandemic, the rigor of analysis was increased through the use of an informatics approach, i.e., Boolean Equivalent Correlated Clusters (BECC^19^; **Figure 1A**) that can identify fundamental invariant (universal) gene expression relationships underlying any biological domain; in this case, the biological domain of ‘*respiratory viral pandemics*’ was selected. Unlike some of the mainstream computational approaches (e.g., differential expression, Bayesian, and correlation network analyses, etc.) that are geared to identify the entire spectra of host immune response, BECC exclusively focuses on Boolean equivalent relationships to identify potentially functionally related gene sets that are part of the invariant spectrum of the host response. The resultant 166-gene ViP signature was found to be conserved in all viral pandemics and outbreaks, and reflected the invariant host immune response to multiple infectious triggers (**Figure 1B** summarizes the types of pathogens that were found to induce the *ViP* signatures^18^). The invariant nature of the host immune response was found to be predominantly *IL15/IL15RA*-centric, and enabled the formulation of precise therapeutic goals and measurement of therapeutic efficacy. At a molecular level, the ViP signatures were distinct from interferon-stimulated genes (ISGs^20,21^), in that, they revealed the broader and fundamental nature of the host immune response, shared between diverse pathogens and tissue/cell types. This included some tell-tale expected (Type I Interferon and cytokine signaling) and some unique (cellular senescence, exhaustion, chromatin silencing, regulation of apoptosis) pathway enrichments^22^. The latter, i.e., the unique pathways, were specifically enriched in a 20-gene subset of the ViP signature, which we called severe (s)ViP signature; this signature was trained on a large dataset of Influenza A/B-infected adult patients annotated with clinical severity^22^. The sViP signature predicted disease severity in COVID-19 (respiratory failure, need for mechanical ventilation, prolonged hospitalization and/or death)^22^. Consequently, the ViP signatures, but not ISGs, were found to be prognostic of disease severity in cohorts of COVID-19 datasets^22^.

**Figure 1:**
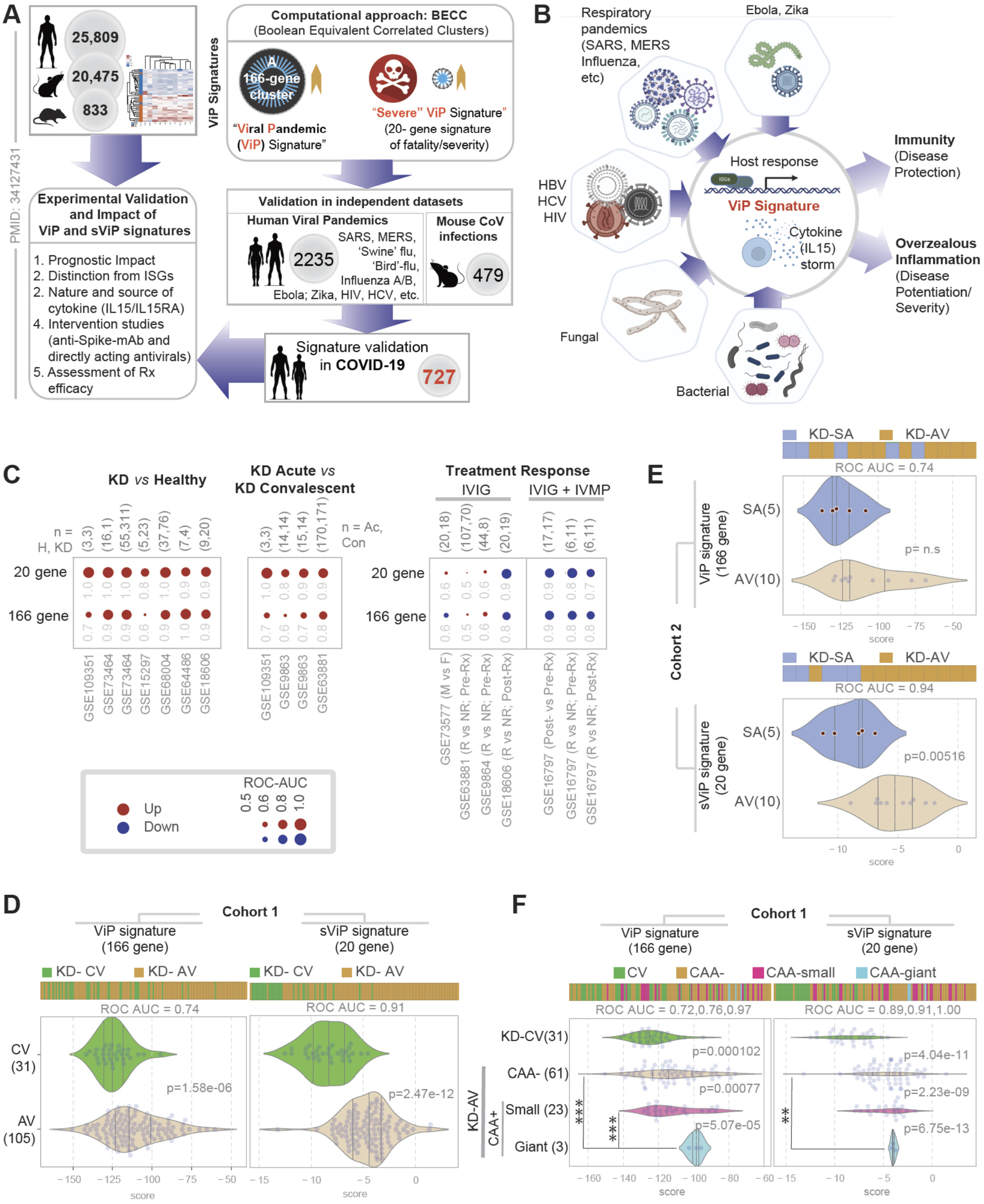
A *<u>Vi</u>ral <u>P</u>*andemic (ViP) signature that is induced in COVID-19^22^, is induced also in epidemic outbreaks of KD. **A**. Schematic displays the computational approach (BECC) and rigor (diversity and number of datasets) used to identify the 166-gene ViP and a subset of 20-gene severe (s)ViP signatures, and the subsequent experimentally validated inferences and impact of the same in a recent study^22^. The numbers in grey circles denote the total number of datasets analyzed in each category. **B**. Schematic displays the various pathogenic triggers that induce ViP signatures (many of which are triggers also for KD) and the prominent induction of *IL15/IL15RA* as an invariant nature of the cytokine storm. **C**. Bubble plots of ROC-AUC values (radii of circles are based on the ROC-AUC) demonstrating the strength of classification and the direction of gene regulation (Up, red; Down, blue) for the classification based on the 20-gene severe ViP signature (top) and 166-gene ViP signature (bottom) in numerous publicly available historic datasets. ViP signatures classified KD *vs*. healthy children (left), acute *vs*. convalescent KD (middle) and treatment response in the setting of combination therapy with IV steroids (MP, methylprednisone) and IV IgG alone (IVIG), but not IVIG alone. Numbers on top of bubble plots indicate number of subjects in each comparison group. **D-E**. Bar (top) and violin (bottom) plots display the classification of blood samples that were collected during acute (AV), sub-acute (SA; ~10-14 d postdischarge) and convalescent (CV; 1 y post-onset) visits from two independent KD cohorts (**D**; Historic Cohort 1**,; E**; Prospective Cohort 2) using ViP (left) or sViP (right) signatures. **F**. Bar (top) and violin (bottom) plots display the sub-classification of blood samples in Cohort 1 based on coronary artery aneurysm (CAA) status using ViP (left) or sViP (right) signatures. Welch’s two sample unpaired t-test is performed on the composite gene signature score to compute the p values. In multi-group setting each group is compared to the first control group and only significant p values are displayed. The p values for comparisons between CAA-vs CAA+ subgroups within acute KD were as follows: CAA-*vs*. CAA-giant, ViP = ***/0.0013, sViP = **/0.033; CAA-small *vs*. CAA-giant, ViP = ***/0.0027, sViP = 0.074; CAA-*vs*. CAA-small, ViP = 0.0087, sViP = 0.099.

Here we used these ViP signatures as is, without further training, as quantitative and qualitative frameworks for measuring the immune response in publicly available historic cohorts of KD predating COVID-19. Both ViP and sViP signatures were upregulated in blood and tissue samples derived from patients with KD compared to healthy controls (ROC AUC for classification of KD vs healthy ranged from 0.8-1.00 across 7 independent cohorts; **Figure 1C**, *left*), and that such induction was limited to the acute phase of KD and downregulated during convalescence (ROC AUC for classification of KD vs healthy ranged from 0.6-0.8 for ViP and 0.8-1.00 for sViP across 4 independent cohorts; **Figure 1C**, middle).

The strength of association between ViP/sViP signatures and acute KD was also preserved in datasets comprised of KD samples prospectively collected before and after IVIG treatment, and treatment response was annotated as responder (R) or non-responder (NR) (**Figure 1C**, right). First, sex had no impact on the induction of signatures (ROC AUC 0.6 in Males vs Females). Second, although the ViP/sViP signatures did not predict treatment response to IVIG (ROC AUC 0.5-0.6 in pre-treatment samples R vs NR; <u>GSE63881</u> and <u>GSE9864</u>), they were reduced in all responders compared to non-responders (ROC AUC 0.8-0.9 in post-treatment samples R vs NR; <u>GSE18606</u>). Finally, in a study^23^ in which the intervention was a combination of IVIG with the intravenous methylprednisolone (IVMP), both ViP signatures were reduced post-Rx (ROC AUC 0.9; <u>GSE16797</u>), and the signatures performed equally well in both pre-treatment and post-treatment samples in differentiating responders from non-responders (ROC AUC 0.7-0.8). These findings suggest that while the IVIG-IVMP combination regimen reduced the signatures effectively among all patients (n = 17), responders induced the ViP signatures to a lesser extent than non-responders. The 20-gene sViP signature consistently outperformed the 166-gene ViP signature in its ability to classify samples across all cohorts tested (**Figure 1C**).

We next confirmed that both the ViP signatures are induced in acute KD (at presentation, < = 10 day of illness) compared to convalescent KD (day 289-3240 of Illness) in a large new cohort of consecutive patients (n=105) who were diagnosed with the disease prior to the onset of the COVID-19 pandemic (Cohort 1; **Supplementary Information 1**) (**Figure 1D**). Again, the sViP signature outperformed the ViP signature in sample classification (ROC AUC 0.91 vs. 0.74). In an independent cohort (Cohort 2, n=20, **Supplementary Information 1; Figure 1E**) prospectively enrolled in the current study after the onset of the COVID-19 pandemic, the ViP signatures could differentiate the acute from subacute (~10-14 d after discharge; ~day 17-25 of Illness) KD samples. As before, the 20-gene sViP signature outperformed the 166-gene ViP signature.

Finally, we tested the association between sViP signatures and markers of disease severity. Because CAA diameter is a predictor of coronary sequelae (thrombosis, stenosis, and obstruction)^24, 25^ and subsequent major adverse cardiac events (unstable angina, myocardial infarction, and death^26^), we used the development of coronary artery aneurysms (CAA) as a marker of disease severity. We found that both ViP signatures differentiated acute KD with giant CAAs (defined as a z-score of ≥10 or a diameter of ≥8 mm^27, 28^) from convalescent KD samples (ROC AUC 0.95 and 0.97 for ViP/sViP signatures, respectively; **Figure 1F**). The ViP signature effectively subclassified acute KD patients with giant aneurysms (CAA-giant) from to those with either no aneurysms (CAA-; p value 0.0027) or small aneurysms (CAA-small; p value p value 0.0013). Similarly, the sViP signature effectively classified acute KD patients with giant aneurysms (CAA-giant) from to those with no aneurysms (CAA-; p value 0.033). Such an analysis was not possible in Cohort 2 (**Supplementary Information 1**) because of the smaller cohort size and absence of subjects with giant CAAs.

We conclude that ViP signatures are induced in acute KD, and track disease severity, i.e., risk of developing giant CAAs, much like we observed previously in the setting of adult COVID-19^22^. Because ViP signatures represent the host immune response to diverse pathogens (**Figure 1B**), upregulation of ViP signatures in KD is consistent with the hypothesis that KD is triggered by multiple infectious triggers ^7, 29, 30^, some of which may be viral in nature^31–33^.

### Comparison of patients with MIS-C and Kawasaki disease

Ten children were included who met the CDC definitions for MIS-C, with detectable anti-SARS-CoV-2 nucleocapsid IgG antibodies [Abbott Architect™] and undetectable virus by polymerase chain reaction (PCR; see **Table 1**). The MIS-C and KD cohorts had notable differences. Although sex and ethnicity were not different, the median age was higher (8.8 years) in the MIS-C cohort than in KD (**Table 1**), which is in keeping with our original report describing this syndrome in June 2020^2^. Left ventricular ejection fraction (LVEF) was reduced in the MIS-C cohort (p = 0.006), consistent with multiple prior reports^34–36^. While all patients had evidence of a marked inflammatory state, the MIS-C cohort had significant cytopenias, including low total WBC, absolute lymphocyte, absolute eosinophil, and platelet counts, with elevation of C-reactive protein level significantly above those observed in the KD cohort (**Table 1**). Most patients (90%) received intravenous immunoglobulin (IVIG) and 70% were treated with intravenous corticosteroids. One patient received anakinra, and three received infliximab. All patients made a full recovery. In all cases, blood was collected for serum before the initiation of any treatments.

**Table 1.**
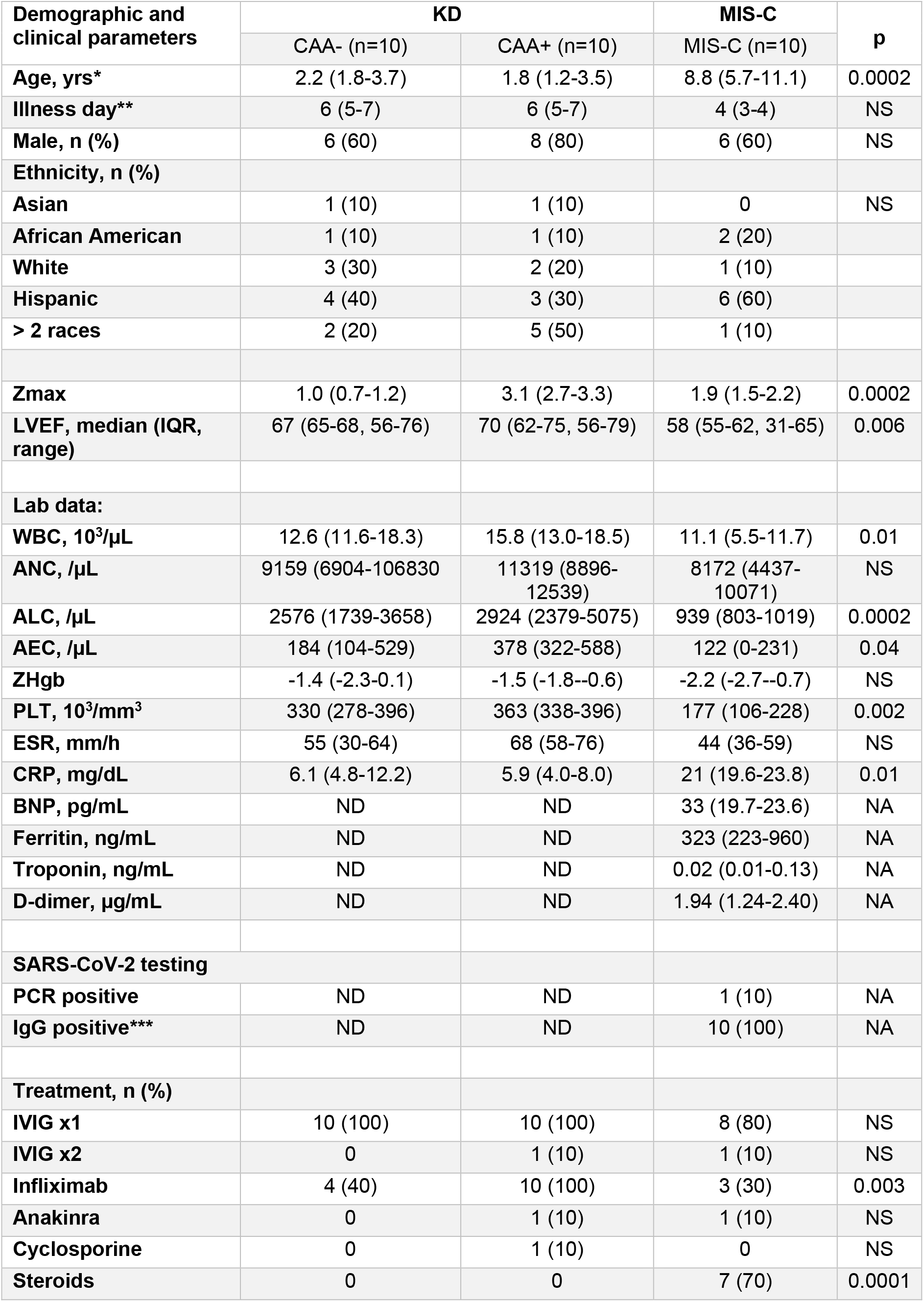
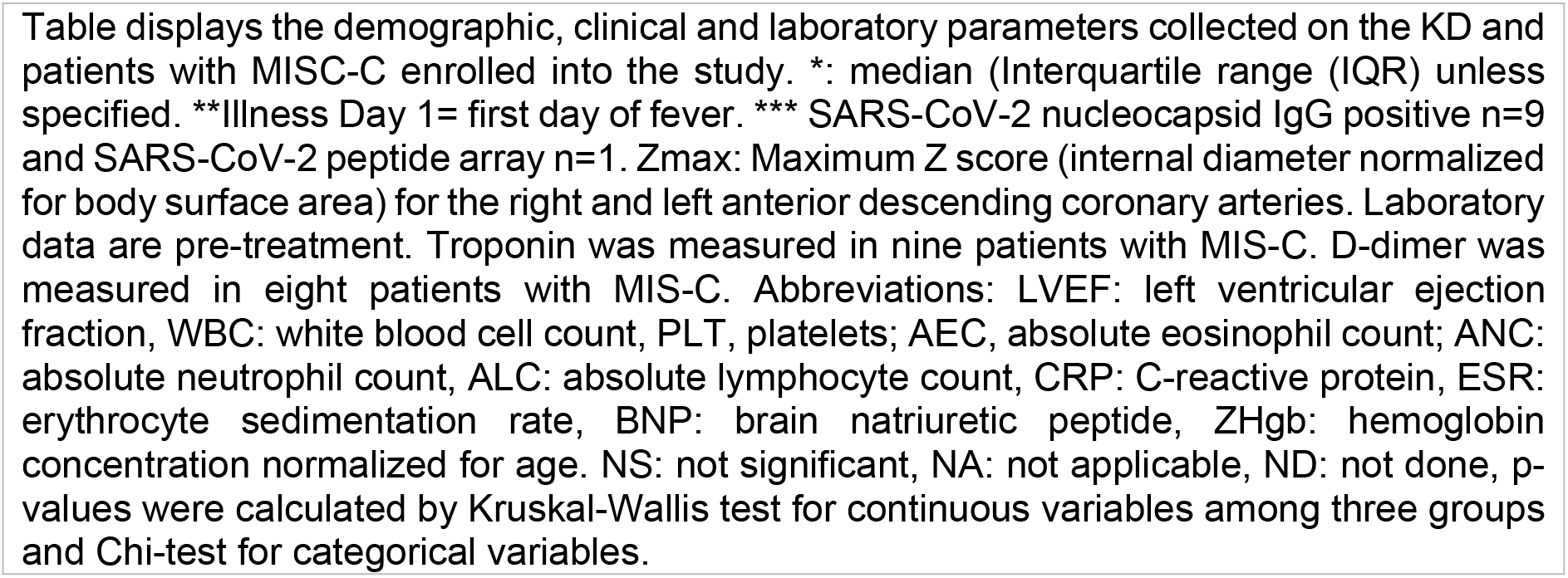
Characteristics of patients with Kawasaki disease (KD) and MIS-C analyzed in this study. Table displays the demographic, clinical and laboratory parameters collected on the KD and patients with MISC-C enrolled into the study. *: median (Interquartile range (IQR) unless specified. **Illness Day 1=first day of fever. *** SARS-CoV-2 nucleocapsid IgG positive n=9 and SARS-CoV-2 peptide array n=1. Zmax: Maximum Z score (internal diameter normalized for body surface area) for the right and left anterior descending coronary arteries. Laboratory data are pre-treatment. Troponin was measured in nine patients with MIS-C. D-dimer was measured in eight patients with MIS-C. Abbreviations: LVEF: left ventricular ejection fraction, WBC: white blood cell count, PLT, platelets; AEC, absolute eosinophil count; ANC: absolute neutrophil count, ALC: absolute lymphocyte count, CRP: C-reactive protein, ESR: erythrocyte sedimentation rate, BNP: brain natriuretic peptide, ZHgb: hemoglobin concentration normalized for age. NS: not significant, NA: not applicable, ND: not done, p-values were calculated by Kruskal-Wallis test for continuous variables among three groups and Chi-test for categorical variables.

| Demographic and clinical parameters | KD |  | MIS-C | P |
| --- | --- | --- | --- | --- |
|  | CAA- (n=10) | CAA+ (n=10) | MIS-C (n=10) |  |
| Age, yrs* | 2.2 (1.8-3.7) | 1.8 (1.2-3.5) | 8.8 (5.7-11.1) | 0.0002 |
| Illness day** | 6 (5-7) | 6 (5-7) | 4 (3-4) | NS |
| Male, n (%) | 6 (60) | 8 (80) | 6 (60) | NS |
| Ethnicity, n (%) |  |  |  |  |
| Asian | 1 (10) | 1 (10) | 0 | NS |
| African American | 1 (10) | 1 (10) | 2 (20) |  |
| White | 3 (30) | 2 (20) | 1 (10) |  |
| Hispanic | 4 (40) | 3 (30) | 6 (60) |  |
| > 2 races | 2 (20) | 5 (50) | 1 (10) |  |
| Zmax | 1.0 (0.7-1.2) | 3.1 (2.7-3.3) | 1.9 (1.5-2.2) | 0.0002 |
| LVEF, median (IQR, range) | 67 (65-68, 56-76) | 70 (62-75, 56-79) | 58 (55-62, 31-65) | 0.006 |
| Lab data: |  |  |  |  |
| WBC, 10<sup>3</sup>/μL | 12.6 (11.6-18.3) | 15.8 (13.0-18.5) | 11.1 (5.5-11.7) | 0.01 |
| ANC, /μL | 9159 (6904-106830) | 11319 (8896-12539) | 8172 (4437-10071) | NS |
| ALC, /μL | 2576 (1739-3658) | 2924 (2379-5075) | 939 (803-1019) | 0.0002 |
| AEC, /μL | 184 (104-529) | 378 (322-588) | 122 (0-231) | 0.04 |
| ZHgb | -1.4 (-2.3-0.1) | -1.5 (-1.8--0.6) | -2.2 (-2.7--0.7) | NS |
| PLT, 10<sup>3</sup>/mm<sup>3</sup> | 330 (278-396) | 363 (338-396) | 177 (106-228) | 0.002 |
| ESR, mm/h | 55 (30-64) | 68 (58-76) | 44 (36-59) | NS |
| CRP, mg/dL | 6.1 (4.8-12.2) | 5.9 (4.0-8.0) | 21 (19.6-23.8) | 0.01 |
| BNP, pg/mL | ND | ND | 33 (19.7-23.6) | NA |
| Ferritin, ng/mL | ND | ND | 323 (223-960) | NA |
| Troponin, ng/mL | ND | ND | 0.02 (0.01-0.13) | NA |
| D-dimer, μg/mL | ND | ND | 1.94 (1.24-2.40) | NA |
| SARS-CoV-2 testing |  |  |  |  |
| PCR positive | ND | ND | 1 (10) | NA |
| IgG positive*** | ND | ND | 10 (100) | NA |
| Treatment, n (%) |  |  |  |  |
| IVIG x1 | 10 (100) | 10 (100) | 8 (80) | NS |
| IVIG x2 | 0 | 1 (10) | 1 (10) | NS |
| Infliximab | 4 (40) | 10 (100) | 3 (30) | 0.003 |
| Anakinra | 0 | 1 (10) | 1 (10) | NS |
| Cyclosporine | 0 | 1 (10) | 0 | NS |
| Steroids | 0 | 0 | 7 (70) | 0.0001 |

### ViP/sViP signatures place MIS-C and KD on the same host immune continuum, but MIS-C as farther along the spectrum than KD

We next analyzed whole blood-derived transcriptome and serum cytokine arrays in the current cohort of subjects with KD (Cohorts 2 and 4) and MIS-C (**Figure 2A**). When MIS-C and acute KD groups were each compared to the control (subacute KD) samples, both ViP (**Figure 2B**) and sViP (**Figure 2C**) signatures were found to be induced at significantly higher levels in MIS-C samples compared to acute KD. However, when MIS-C and acute KD were compared to each other, we found that the ViP signatures could not distinguish between these samples, indicating that both conditions share a similar host immune response. Heatmaps of patterns of expression (**Figure 2D-E**) demonstrate that most of the individual genes contributed to the elevated ViP and sViP signatures observed in MIS-C samples. These genes included *IL15* and *IL15RA* (highlighted in red; **Figure 2D**), both components within an invariant cytokine pathway that was previously demonstrated to be elevated in the lungs of patients with fatal COVID-19 and in SARS-CoV-2 challenged hamsters^22^.

**Figure 2.**
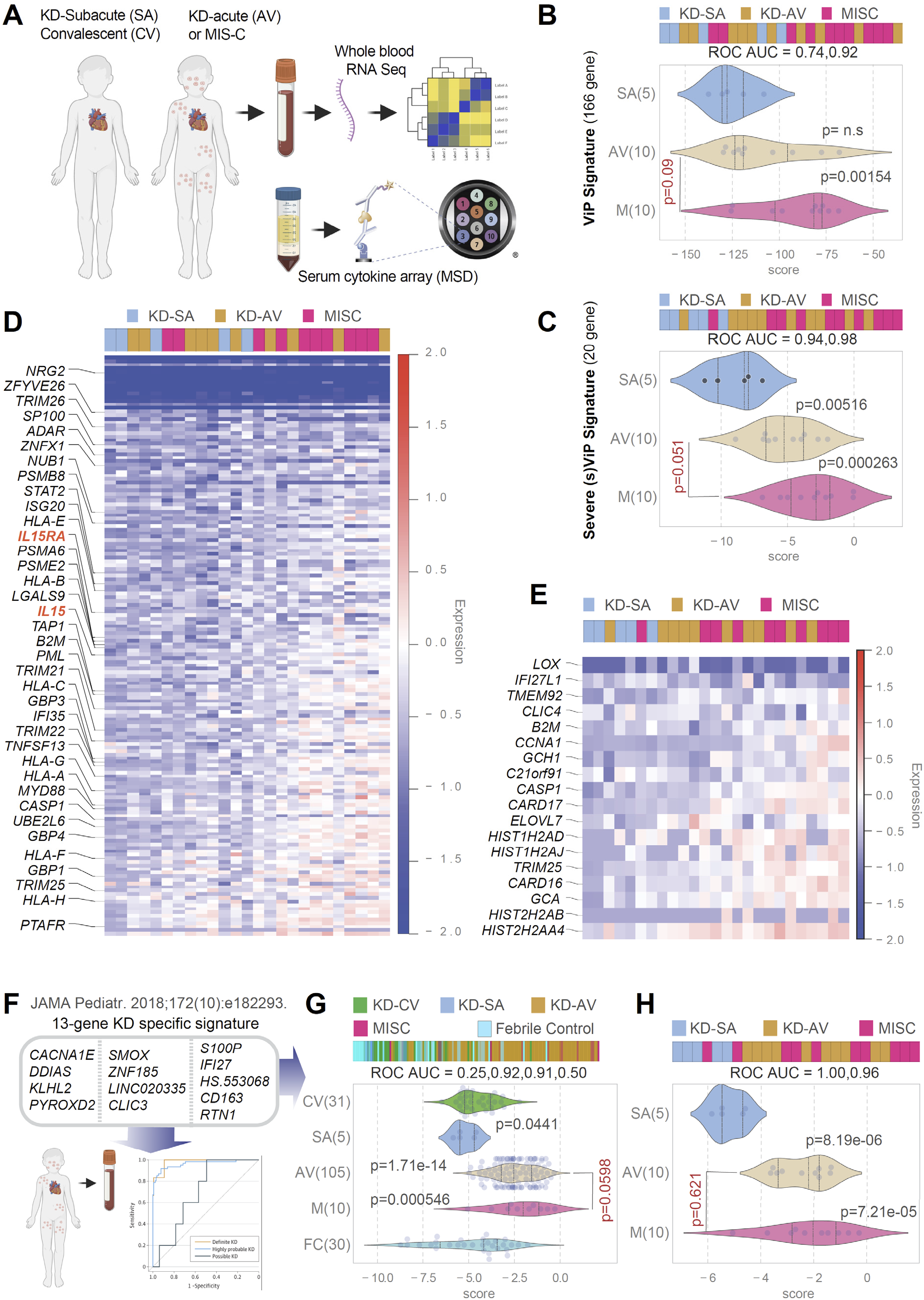
A KD-specific 13 transcript signature^38^ shows that KD and MIS-C are indistinguishable, but ViP/sViP signatures place MIS-C as farther along the spectrum than KD. **A**. Schematic displays the workflow for patient blood collection and analysis by RNA Seq (this figure) and cytokine array by mesoscale (**Figure 4-5**). **B-C**. Bar (top) and violin (bottom) plots display the classification of blood samples that were collected during collected during acute (AV) and sub-acute (SA; ~10-14 d post-discharge) visits of KD subjects and from patients diagnosed with MIS-C. The p value for comparison between acute KD (AV) and MIS-C (M) is displayed in red font. **D-E**. Heatmaps display the patterns of expression of the 166 genes in ViP (D) and 20 gene sViP (E) signatures in the KD and MIS-C samples. The only cytokine-receptor pair within the signature, i.e., *IL15/IL15RA*, are highlighted on the left in red font in D. **F**. Schematic displays the 13-transcript whole blood signature (no overlaps with ViP signature genes) previously demonstrated to distinguish KD from other childhood febrile illnesses^38^. **G-H**. Bar (top) and violin (bottom) plots display the classification of blood samples that were collected during acute (AV) and convalescent (CV) visits from two independent KD cohorts (**G**; Historic Cohort 1**; E**; Prospective Cohort 2) using 13-transcript KD signature. FC, febrile control. See also **Supplementary fig. S1** for co-dependence analysis of ViP and KD-13 signatures. Welch’s two sample unpaired t-test is performed on the composite gene signature score to compute the p values. In multi-group setting each group is compared to the first control group and only significant p values are displayed. The p value for comparison between acute KD (AV) and MIS-C (M) is displayed in red font.

Taken together, these analyses led to two key conclusions: (i) that the host immune response, as detected in a qualitative manner using the ViP signatures, is similar in KD and MIS-C and has a *IL15/IL15RA* invariant component; (ii) that the degree of such host immune response, as measured quantitatively using the ViP signature scores, is more intense in MIS-C than KD. These findings are consistent with the fact that MIS-C is a host immune response to SARS-CoV-2 exposure, and we previously showed that the interaction of viral spike protein with the host entry receptor, ACE2 is critical for the induction of ViP signatures^22^. Findings are also in keeping with prior work^37^ showing that serum levels of IL15 is significantly elevated in acute KD, ~10-fold compared with subacute-KD and normal controls, and that such increase correlated with the concomitant increase in serum TNFα.

### A KD-specific signature independently confirms that KD and MIS-C are syndromes on the same host immune response continuum

To circumvent an over-reliance on one set of signatures (i.e., ViP/sViP), we next analyzed a KD-specific 13 transcript diagnostic signature^38^ that was previously shown to be effective in distinguishing children with KD from all other febrile conditions. During validation, the 13-transcript signature mirrored the certainty of clinical diagnosis, i.e., it differentiated definite, highly probable, and possible KD from non-KD with ROC AUCs of 98.1% (95% CI, 94.5%-100%), 96.3% (95% CI, 93.3%-99.4%), and 70.0% (95% CI, 53.4%-86.6%), respectively (**Figure 2F**). Unlike the ViP signatures, which has a typical enrichment of interferon and cytokine pathways with a prominent presence of *IL15/IL15RA*, the KD-signature is comprised of a set of non-overlapping genes, some of which relate to major central hubs within the tumor necrosis factor (TNFα) and interleukin 6 (IL6) pathways^38^. When we applied this signature to the historic cohort 1 (**Figure 2G**) and to our current cohort (Cohort 2; **Figure 2H**), we found that the KD-specific 13 transcript signature could not distinguish between MIS-C and KD in either cohort. Furthermore, a correlation test demonstrated that the two non-overlapping signatures, sViP and KD-13, both of which are significantly induced in KD and MIS-C (**Figure 2C**) are independent of each other (**Supplementary fig. S1**). This suggests that these two signatures reflect two fundamentally distinct and unrelated biological domains within the host immune response; whether their diagnostic/prognostic abilities may have an additive benefit remains to be explored.

The similar extent to which KD and MIS-C induced the KD-13 signature in two independent cohorts further supports our observation with ViP/sViP signatures that KD and MIS-C share fundamental aspects of host immune response with each other. That KD and MIS-C samples share ViP/sViP signatures with COVID-19 implies that the three diseases represent distinct clinical states on the same host immune response continuum.

### The sViP signature can recognize severe form of MIS-C that presents with myocardial dysfunction

Next, we asked if the sViP signature can track disease severity in MIS-C. Because of the limited number of ‘severe’ cases in our MIS-C cohort, we prospectively analyzed two recently accessible MIS-C cohorts (<u>GSE166489</u> ^15^ and <u>GSE167028</u> ^39^). While both datasets analyzed PBMCs from MIS-C subjects, and both studies used the presence of myocardial dysfunction as basis for severe disease, each study used a slightly different criterion for classification of disease severity (**Figure 3A**). de Cevins et al.,^39^ classified MIS-C as severe when the patients presented with elevated cardiac troponin I and/or altered ventricular contractility by echocardiography, and clinical signs of heart failure requiring ICU support. Ramaswamy et al.,^15^ classified MIS-C as severe if they were critically ill, with cardiac and/or pulmonary failure. In both cohorts, sViP was able to classify severe MIS-C (with myocardial dysfunction; MYO+) from mild-moderate disease (who recovered or presented without myocardial dysfunction; MYO-) (**Figure 3B-C**); while the p value was significant in <u>GSE166489</u> (**Figure 3B**), a similar trend was conserved in <u>GSE167028</u> ^39^ (**Figure 3C**). These findings show that the sViP signature can identify severe MIS-C who are at risk to develop myocardial dysfunction, just as it did in the case of KD subjects who are at risk of developing giant CAAs (**Figure 1F**) and similar to its prior performance in identifying adults with COVID-19 who are at risk of respiratory failure, mechanical ventilation, prolonged hospitalization and/or death^22^.

**Figure 3:**
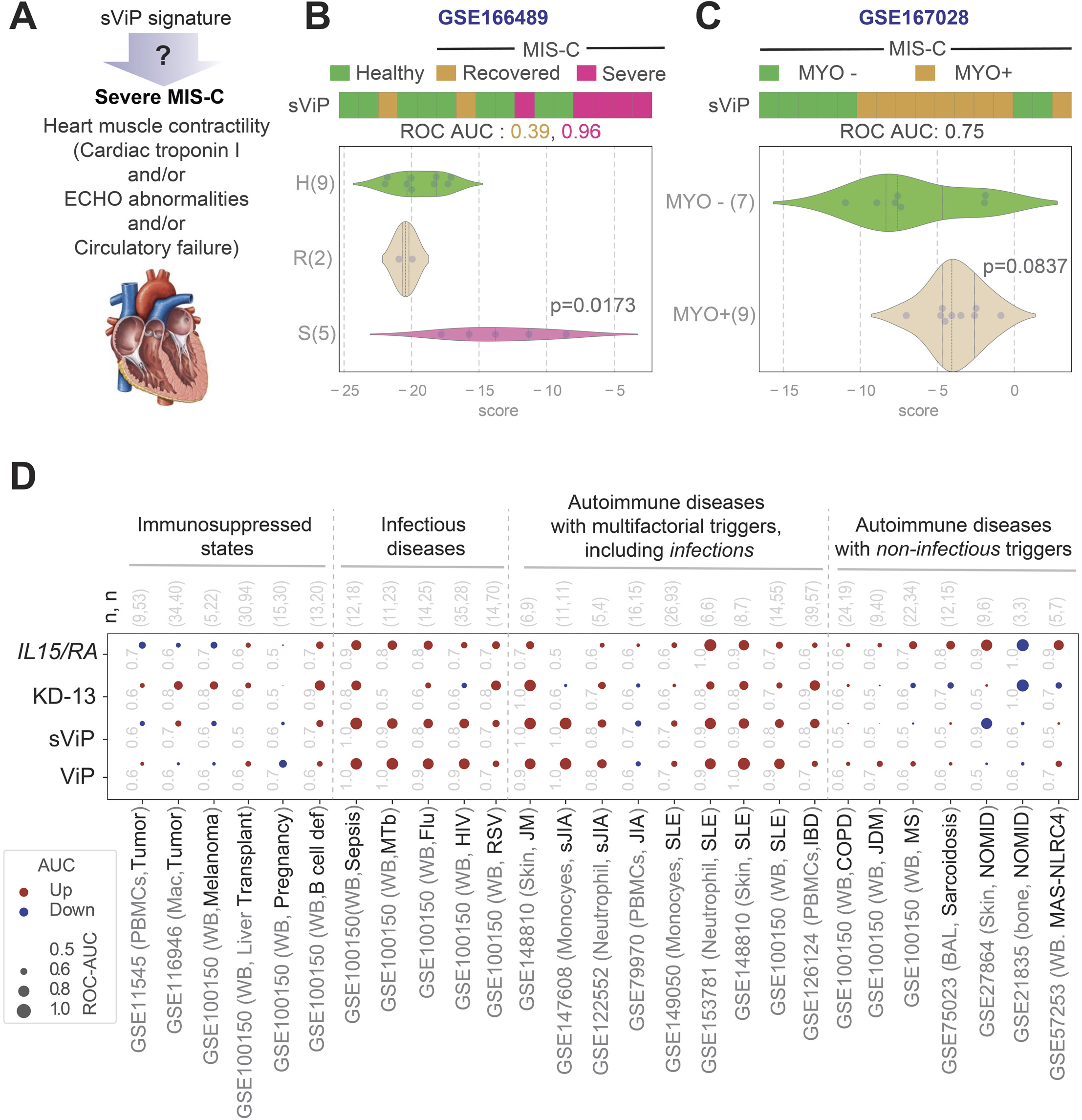
Performance of ViP/sViP signatures on independent MIS-C datasets and on diverse tissues and in diverse diseases of the immune system. **A-C**. Severe (s)ViP signature can classify severe MIS-C based on transcriptomic analysis of blood that was prospectively collected at admission in two independent studies (<u>GSE166489</u> ^15^ and <u>GSE167028</u> ^39^). Schematic in **A** summarizes the definition of severe MIS-C used by two independent studies. **B-C**. Bar (top) and violin (bottom) plots display the classification of blood samples in two cohorts of MIS-C subjects, based on the need for ICU management due to the presence (MYO+) or recovery in the absence (R or MYO-) of myocardial dysfunction using sViP signature. Welch’s two sample unpaired t-test is performed on the composite gene signature score to compute the p values. **D**. Bubble plots of ROC-AUC values (radii of circles are based on the ROC-AUC) demonstrating the strength of classification and the direction of gene regulation (Up, red; Down, blue) in numerous publicly available historic datasets representing diverse diseases of the immune system using 4 gene signatures: the 166-gene ViP signature, the 20-gene sViP signature, the KD-13 signature, and finally the *IL15/IL15RA* composite score. Numbers on top of bubble plots indicate number (n) of control *vs*. disease samples in each dataset. Abbreviations: PBMCs, peripheral blood mononuclear cells; Mac, macrophages; WB, whole blood; MTb, *M. tubercutosis*; Flu, Influenza; HIV, Human immunodeficiency virus; RSV, Respiratory syncytial virus; JM, Juvenile myositis; sJIA, Systemic Juvenile Idiopathic Arthritis; SLE, Systemic lupus erythematosus; IBD, Inflammatory bowel disease; COPD, Chronic obstructive pulmonary disesase; JDM, Juvenile dermatomyositis; MS, Multiple sclerosis; BAL, Bronchoalveolar lavage; NOMID, Neonatal onset multisystem inflammatory disease; MAS, Macrophage activation syndrome; NLRC4, NLR Family CARD Domain Containing 4.

Taken together with the prior findings, we conclude that the 20-gene sViP signature captures a core set of genes that are expressed in the setting of an overzealous (prolonged or intense, or both) host immune response in all three diseases— KD, MIS-C (this work) and COVID-19^22^ – despite the fact that each present with distinct clinical features of severity.

Because all three conditions represent diseases of the immune system that share an ‘infectious trigger’, we asked if the ViP/sViP signatures are also induced in the setting of other diseases of the immune system. To this end, we analyzed numerous publicly available datasets, ranging from immunosuppressed states (as negative control), infectious diseases (both viral and bacterial; as positive control), and autoimmune diseases (**Figure 3D**), and assessed the ability of ViP/sViP signatures to classify control and diseased samples in each dataset. Because the ViP/sViP signatures are able to detect the shared core fundamental host immune response in cell/tissue agnostic manner^22^, we tested diverse samples ranging from whole blood to bronchoalveolar lavage fluid (**Figure 3D**). The ViP/sViP signatures performed as anticipated in the negative and positive control datasets, i.e., neither signature was induced in immunosuppressed conditions, e.g., malignancies, pregnancy, posttransplant immunosuppression, but both were induced in infectious diseases, e.g., sepsis, HIV, RSV, and tuberculosis (*left*; **Figure 3D**). In the case of the autoimmune diseases, the ViP/sViP signatures were induced in some, but not others. The signatures were induced in those conditions that have multifactorial triggers, including potential contributions from infections; for example, mechanistic studies have identified viral link in many of them (EBV-linked autoimmune diseases^40^ such as JIA, SLE, IBD). The signatures were not induced in other conditions where the disease triggers remain mysterious (e.g., sarcoidosis) or where the disease is driven by specific mutations, e.g., Neonatal onset multisystem inflammatory disease (NOMID) that is due to mutant NLRC3 and macrophage activation syndrome (MAS) that is due to mutant NLRC4). These findings lend further support to our finding that ViP/sViP signatures are induced and perform well to identify severe MIS-C, which shares infection as a trigger, much like KD and COVID-19. Intriguingly, numerous infectious and autoimmune diseases shared the *IL15/IL15RA*-centric cytokine response, which is in keeping with prior observations^41^.

### Cytokine panels and whole blood transcriptomes reveal subtle differences between MIS-C and KD

We next analyzed a set of 10 serum cytokines using Meso Scale Discovery Electrochemiluminescence (MSD-ECL) Ultra-Sensitive Biomarker Assay. A panel of 10 target cytokines was prioritized based on a review of the literature for the reported presence and/or relevance of each in either KD and/or MIS-C. An unsupervised clustering of just these 10 cytokines was sufficient to differentiate acute KD and MIS-C from one-year convalescent KD samples (**Figure 4A**; Cohorts #2 and #3, **Supplementary Information 1**); the convalescent samples served as baseline ‘healthy’ controls in this case. Regardless of their degree of elevation in the acute setting, all cytokines were virtually undetectable in convalescent samples (**Supplementary fig. S2**). While most cytokines were induced indistinguishably in acute KD and MIS-C (**Figure 4A**; **Figure 4B**, *top*), notable exceptions were TNFα, IFNγ, IL10, IL8 and IL1β, all of which were elevated to a greater extent in MIS-C compared to KD (**Figure 4B**, *bottom*), either significantly (TNFα, IFNγ; **Figure 4B**) or trended similarly, but failed to reach statistical significance (IL10, IL1β, IL8). Gene set enrichment analyses (GSEA) on the transcriptomic dataset for each of the differentially expressed cytokines (**Figure 4C**) showed that the gene sets for those pathways were also induced in MIS-C at levels significantly higher than KD (**Figure 4D-H**).

**Figure 4:**
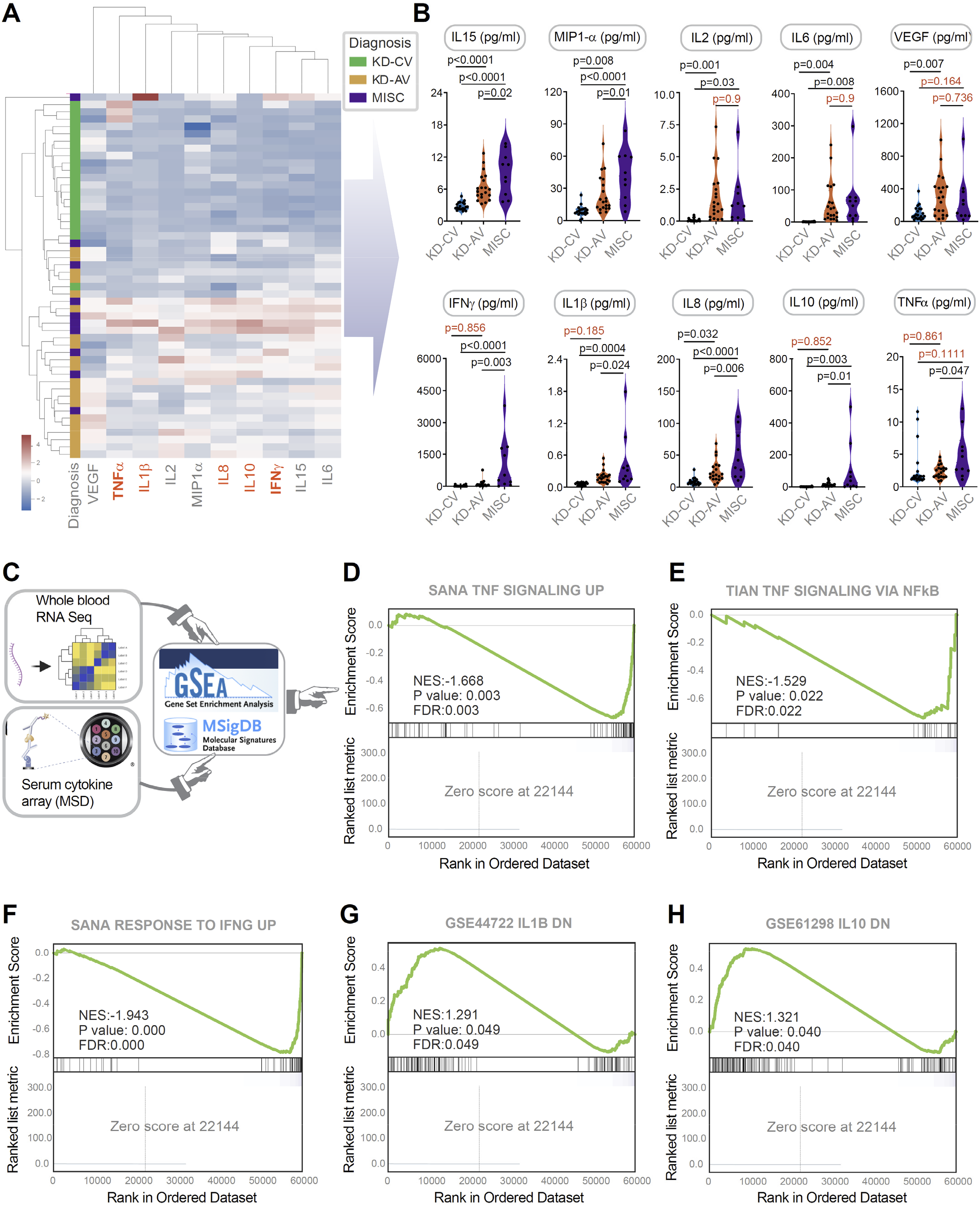
Serum cytokine arrays and whole blood transcriptomes reveal the severity and nature of the cytokine storm in MIS-C that distinguishes it from KD. **A**. Heatmap displays the results of unsupervised clustering of sub-acute and acute KD (KD-SA, KD-AV; n = 10 each) and MIS-C (n = 10) subjects using the cytokine profiles determined by mesoscale (MSD). Red = cytokines differentially expressed between MIS-C and KD. See also **Supplementary fig. S2** for violin plots for individual cytokines. **B**. Violin plots display the shared (top panels; IL15, MIP1a, IL2, IL6 and VEGF) and distinct (bottom panels; IFNγ, IL1β, IL8, IL10 and TNFα) features of the cytokine storm in MIS-C vs KD subjects. Statistical significance was determined by one-way ANOVA followed by Tukey’s test for multiple comparisons. **C**. Schematic shows the process used to integrate serum cytokine array results with whole blood RNA Seq data; cytokines that were differentially expressed in MIS-C were used to inform GSEA of the corresponding pathways. **D-F**. Gene Set Enrichment Analysis (GSEA preranked analysis) of three pathways derived from MSigDB: SANA TNF SIGNALING UP (D), TIAN TNF SIGNALING VIA NFkB (E), and SANA RESPONSE TO IFNG UP (F) demonstrate the significance of TNF (D-E) and IFNG (F) pathway activation in MIS-C. **G-H**. Down-regulated genes after IL1B (G) and IL10 (H) stimulation were derived from differential expression analysis of GSE44722 (n = 269 genes), and GSE61298 (n = 208 genes) respectively. GSEA pre-ranked analysis to test the significance of IL1B and IL10 pathway is performed like panel D-F using the down-regulated genes. Source data are provided as a Source Data file.

Taken together, these findings suggest that FDA-approved therapeutics targeting TNFα and IL1β pathways may be beneficial to treat MIS-C. The IL-1 receptor is expressed in nearly all tissues and its antagonism by anakinra, a recombinant form of IL-1Ra^42^, prevents receptor binding of either IL-1α or IL-1β. Similarly, infliximab, a chimeric antibody to TNFα, has been repurposed for COVID-19^43–45^, and our analyses suggest that this agent holds promise as a treatment for MIS-C.

### Integrated analyses of ViP/sViP signatures, cytokine profile, and clinical laboratory parameters reveal unique features of MIS-C and indicators of disease severity

We next sought to understand how similar host cytokine responses can trigger two distinct clinical syndromes, and how such responses may drive features of clinical severity. To this end, we first carried out an agglomerative hierarchical clustering of the MIS-C and acute KD samples using both cytokine profiling (MSD) and clinical/laboratory parameters. This analysis, coupled with correlation tests (**Supplementary fig. S3**) revealed several intriguing observations: (i) visualization by heatmap showed that compared to KD, MIS-C patients had higher cytokine levels and more severe pancytopenia (**Figure 5A**); (ii) although platelet counts (PLT), but not absolute eosinophil counts (AEC) were significantly reduced in MIS-C compared to KD (**Figure 5B**), there was a strong positive correlation between PLT and AEC in MIS-C, but not KD (**Figure 5C**; left) and strong negative correlations of PLT with IL15 in both KD and MIS-C (**Figure 5C**; right) and with MIP1α in MIS-C, but not KD (**Figure 5D**); (iii) this is consistent with the fact that IL15 and MIP1α were found to have a strong positive correlation in MIS-C, but not KD (**Figure 5E**). These findings suggest that MIS-C has key distinguishing features of thrombocytopenia and low eosinophil counts, and that both features are negatively correlated with the serum levels of IL15, a key invariant feature of the ViP signature. These findings also held true when we analyzed the two clinical parameters, PLT and AEC, against ViP/sViP signatures, as well as a specific *IL15/IL15RA* composite transcript score from whole blood RNA Seq dataset. We found that PLT and AEC negatively correlated with ViP (**Figure 5F**; top), sViP (**Figure 5F**; middle) and a *IL15/IL15RA* composite score (**Figure 5G-F**; bottom) in MIS-C, but such correlations were restricted only to PLT in acute KD. These findings indicate that MIS-C, but not KD, has at least two distinct and interrelated clinical features, thrombocytopenia and eosinopenia, that appear to be related to the degree of induction of ViP signatures and a IL15-predominant cytokine induction. Findings also suggest that MIP1α is a key contributor to the immune response in MIS-C and that its levels are closely and positively related to the levels of IL15.

**Figure 5:**
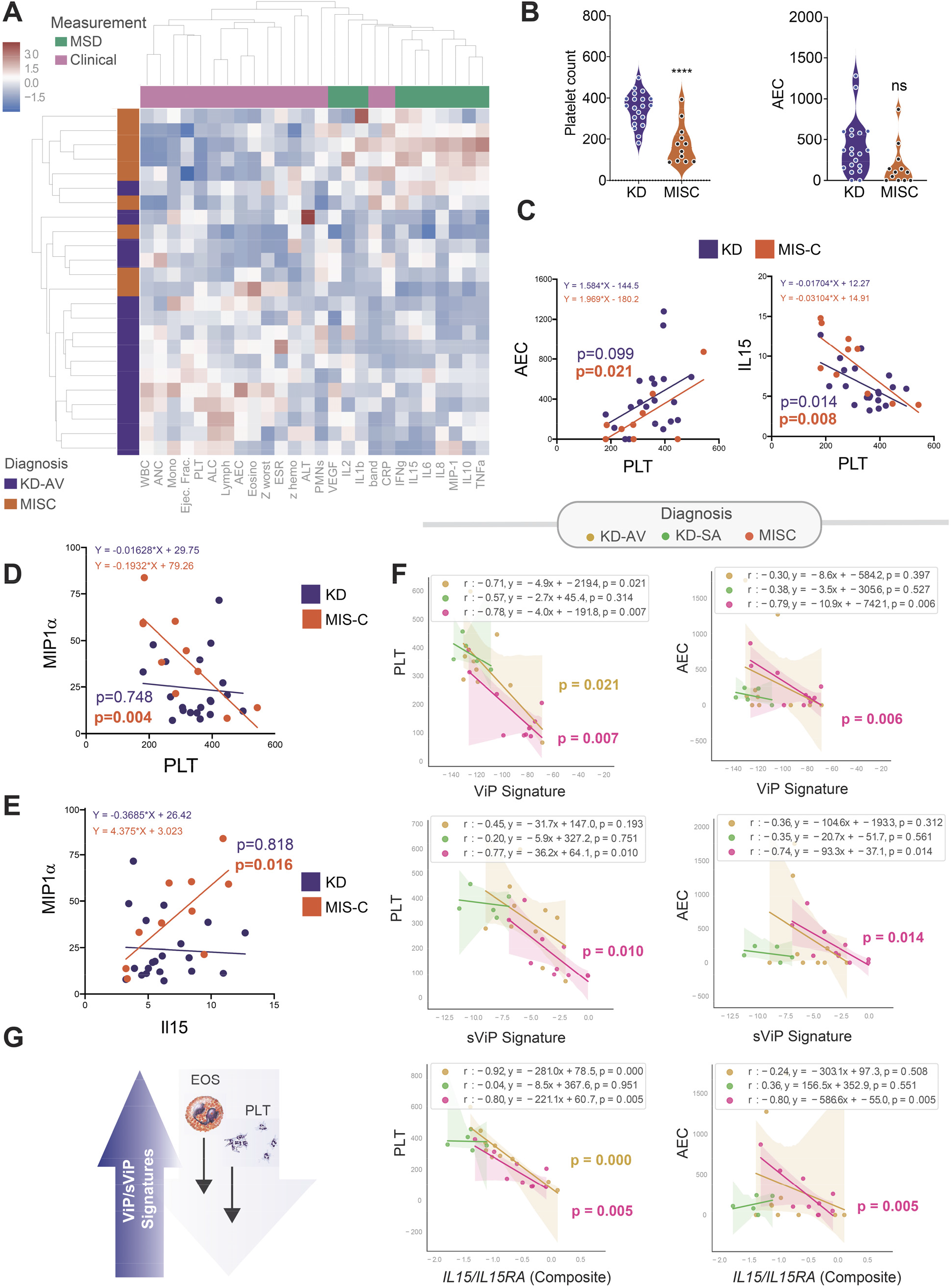
An integrated analysis of mesoscale (cytokine) data, ViP/sViP transcriptomic signatures and laboratory and clinical parameters reveals features that are unique to MIS-C. **A**. Heatmap displays the results of hierarchical agglomerative clustering of acute KD (KD-AV; n = 10) and MIS-C (n = 10) subjects using the cytokine profiles determined by mesoscale (MSD) and the laboratory features. **B**. Violin plots display PLT (platelet) and AEC (absolute eosinophil counts) in KD and MIS-C (unpaired Student’s t-test used to test significance). **C-E**. Correlation test between AEC and PLT (C; left) and IL15 and PLT (C; right), and MIP1α and PLT (D) and MIP1 α and IL15 (E) are shown, and significance was calculated and displayed using GraphPad Prism 9. Significance: ns: non-significant, ****, p < 0.0001. See **Supplementary fig. S3** for all possible correlation tests between clinical and cytokine data in KD, MIS-C and COVID-19. **F**. Correlation tests between PLT (left) or AEC (right) on the Y axis and gene signature scores on the X axis [either ViP (top), sViP (middle) or a *IL15/IL15RA* composite (bottom)] were calculated and displayed as scatter plots using python seaborn lmplots with the p-values. The confidence interval around the regression line is indicated with shades. **G**. Schematic summarizing the findings in MIS-C based on laboratory and RNA seq analysis.

These findings reveal key similarities and differences among MIS-C, KD and COVID-19. Thrombocytopenia, which was more pronounced in MIS-C and correlated significantly with IL15 and *IL15/IL15RA* composite transcript score in both KD and MIS-C, has also been reported in COVID-19 and postulated because of various mechanisms^46–50^. In the case of KD, thrombocytopenia has been found to be associated with disease severity^51^. Similarly, in the case of COVID-19, a large meta-analysis confirmed that approximately 12% of hospitalized patients have thrombocytopenia, which represents a sign of disease severity and poor outcomes^46^. Thrombocytopenia carried a 3-fold enhanced risk of a composite outcome of intensive care unit admission, progression to acute respiratory distress syndrome, and mortality (odds ratio [OR], 3.49; 95% CI, 1.57−7.78), and a subgroup analysis confirmed a significant association with mortality (OR, 7.37; 95% CI, 2.08−26.14). Eosinopenia appears to be a notable shared feature between MIS-C and COVID-19^52^, but not KD. These findings are consistent with the fact that KD is known to present with higher (not lower) eosinophil counts, Th2 cytokines IL-4, IL-5, and eosinophil cationic protein (ECP)^53–57^. As in the case of thrombocytopenia, persistent eosinopenia after admission correlated with COVID-19 severity and low rates of recovery^58^.

### ViP/sViP signatures track the severity of two distinct cardiac phenotypes in MIS-C and KD

We next analyzed the relationship between ViP signatures and the prominent and unique cardiac phenotype in MIS-C reported by others^34–36^ and observed also in our cohort (**Figure 6A**), i.e., a significantly reduced LVEF that can present with cardiogenic shock necessitating ionotropic support. We found that sViP signature scores, but not ViP or *IL15/IL15RA* composite scores correlate significantly with LVEF (**Figure 6B-D**), indicating that LVEF may belong to the domain of clinical indicators of disease severity in MIS-C (alongside platelets and AEC), but it may not be directly related to the *IL15*-centric cytokine signaling. In KD, the ViP and sViP signatures were tested earlier (**Figure 1F**) and found to distinguish patients with giant CAA from convalescent samples with ROC AUC > 0.95. A *IL15/IL15RA* composite score performed similarly in distinguishing those samples (**Figure 6E**). We hypothesized that the ViP signatures may be related to two distinct cardiac phenotypes in severe disease: the signatures in KD may signify the nature of the vasculitis that drives the formation of CAAs, whereas the same signature in MIS-C may signify the degree of cardiomyopathy that impairs contractility (**Figure 6F**). Because we were unable to acquire cardiac tissues from MIS-C-related autopsies, we carried out immunohistochemical analyses on cardiac tissues from a case of fatal KD. We found that both IL15 and IL15RA were expressed in the cardiomyocytes and coronary arterioles amidst extensive fibrosis, as detected using Masson’s trichrome stain (**Figure 6G**).

**Figure 6:**
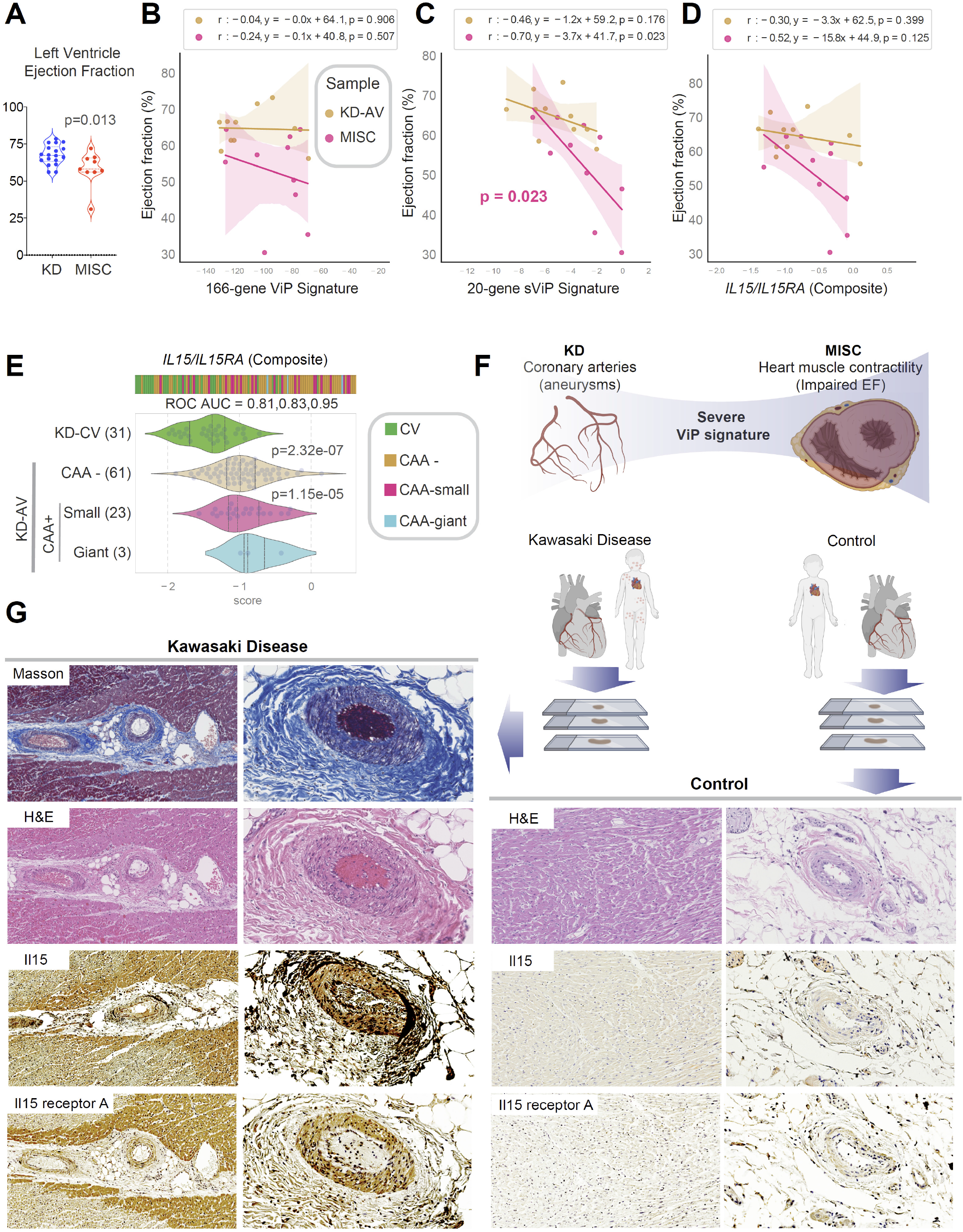
ViP/sViP signatures correlate with two distinct cardiac phenotypes in MIS-C and KD. **A**. Violin plots display the left ventricular ejection functions (LVEF) in KD and MIS-C patients. Statistical significance was determined by unpaired Student’s t-test. **B-D**. Correlation tests between LVEF (Y axis) and gene signature scores on the X axis [either ViP (B), sViP (C) or a *IL15/IL15RA* composite (D)] are displayed as a scatter plot and significance was calculated and displayed as in **5F**. The confidence interval around the regression line is indicated with shades. **E**. Bar and violin plots show how a *IL15/IL15RA* compositive score varies between KD samples. The score classifies KD-AV with giant CAAs from control (KD-CV) samples with a ROC AUC 0.95. Welch’s two sample unpaired t-test is performed on the composite gene signature score to compute the p values. In multi-group setting each group is compared to the first control group and only significant p values are displayed. **F**. Schematic summarizes how sViP signature classifies two distinct cardiac phenotypes, CAA and reduced pump function in KD and MIS-C, respectively. **G**. Consecutive sections from formalin fixed paraffin-embedded (FFPE) cardiac tissue collected at autopsy from a child with severe debilitating cardiac complications of KD were stained for masson’s trichrome, H&E and IL15/IL15 receptor A antigens (left). Healthy heart sections from another child were used as controls (right).

Together, these findings suggest that the *IL15/IL15RA* induction we see in COVID-19, KD and MIS-C may have distinct sources and/or target end organs: We previously showed prominent induction of *IL15/IL15RA* in the lung alveoli of fatal COVID-19 patients^22^, and here we show it in the coronary arteries and cardiomyocytes in KD. Because cardiomyopathy and enteropathy are the commonest presenting features of MIS-C, it is tempting to speculate that cardiac muscle and various regions of the gastrointestinal tract may be the site of *IL15/IL15RA* induction in MIS-C.

## DISCUSSION

Using a combination of publicly available KD datasets and newly recruited cohorts of KD and MIS-C subjects (summarized in **Figure 7A**) and a set of gene signatures we report an unexpected discovery regarding the host immune response in these diagnoses. Our findings show that two distinct clinical syndromes, KD, which predates the current pandemic by 6 decades, and the novel COVID-19, share a similar profile of host immune response. The same host immune response is seen also in MIS-C, a new disease that co-emerged with COVID-19, which has some overlapping features with KD (i.e., clinical presentation, pediatric, etc.), and yet is an immune response to the virus that causes COVID-19 (**Figure 7B**). We assessed the quality and intensity of the host immune response in these syndromes with a powerful and unbiased computational tool, the ViP signatures^22^. Previously we had demonstrated the usefulness of the ViP signatures to define and measure the host immune response in COVID-19, identify the site/source of the cytokine storm, track disease severity, objectively formulate therapeutic goals and track the effectiveness of emerging drugs/biologics^22^. We now show that the same ViP signatures can objectively demonstrate the shared immunophenotypes between all three syndromes, which features an invariant upregulation of the *IL15/IL15RA* pathway. That a 13 transcript KD-specific signature that was previously shown to distinguish KD from other non-KD febrile illnesses^38^ failed to distinguish KD from MIS-C, further confirmed that KD and MIS-C share identical molecular markers of disease and hence, are fundamentally similar at the molecular level (summarized in **Figure 7B**). These findings are in keeping with what has been observed by Consiglio et al.,^13^ who found KD and MIS-C to be clustered together in a PCA analysis of plasma proteins. These findings suggest that the two clinical syndromes not just share common clinical features (e.g., rash, fever, etc.), but may also share proximal pathways of immunopathogenesis.

**Figure 7.**
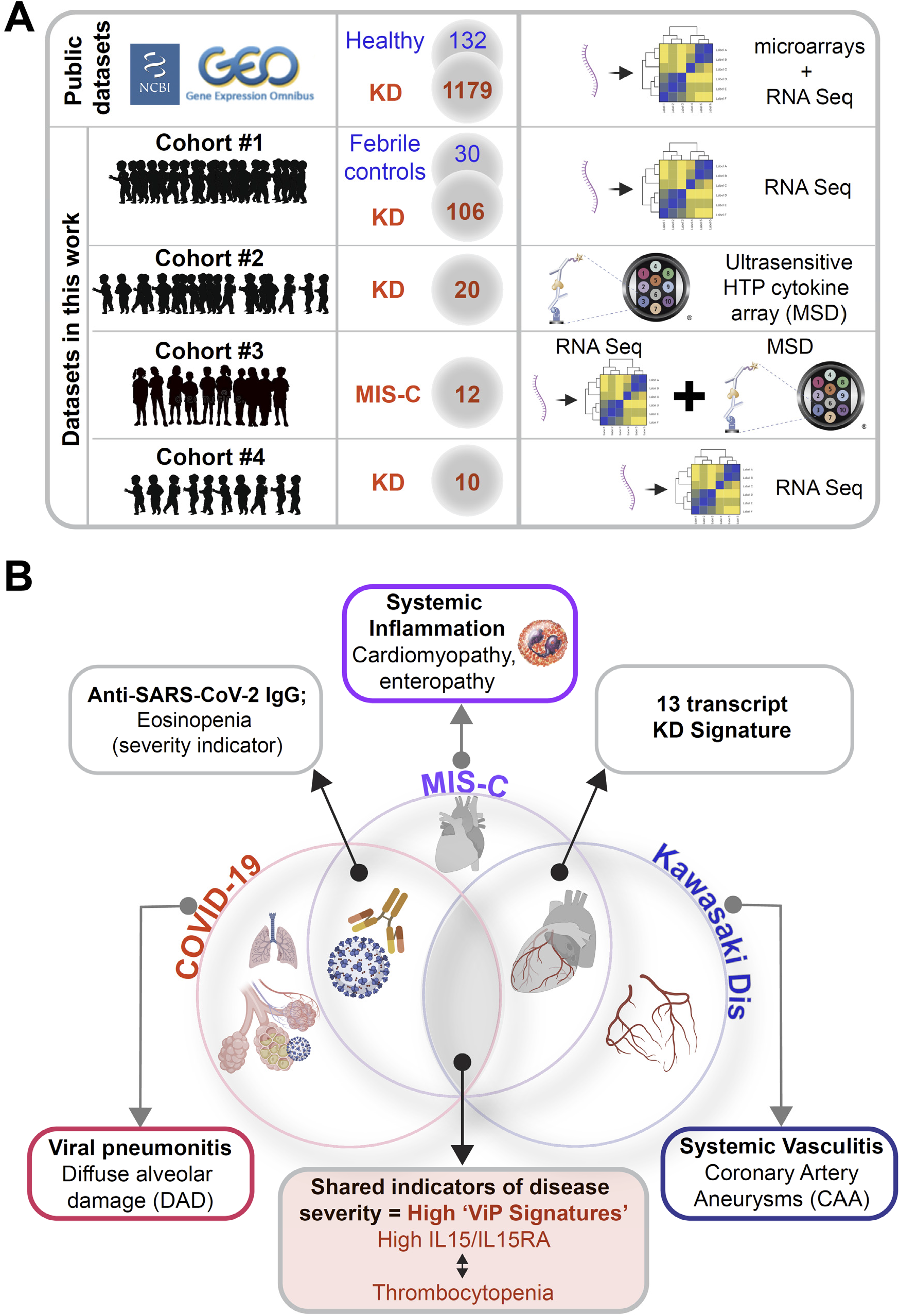
Summary of findings and conclusions. **A.** Summary of datasets used (publicly available prior ones and new original cohorts) to support the conclusions in this work. Numbers in circles denote the number of subjects in each cohort. **B.** Venn diagram displays the major findings from the current work. ViP/sViP signatures, and more specifically, the *IL15/IL15RA* specific gene induction are shared between patients in all three diagnostic groups. While these signatures are known to be associated with diffuse alveolar damage in the lungs of patients with COVID-19^22^, it is associated with CAA in KD and with reduction in cardiac muscle contractility in MIS-C. Overlapping features between each entity are displayed.

Despite the high-level broad similarities, the ViP signatures also helped identify key differences in clinical/laboratory parameters that may help distinguish MIS-C from KD. First, although the ViP signatures placed KD and MIS-C on the same host immune continuum, the degree of host immune response in MIS-C is significantly higher than KD by all measures tested (i.e., ViP, sViP, *IL15/IL15RA* and KD-13 signatures and direct measurement of serum cytokines). Higher ViP signatures in MIS-C tracked three major clinical and/or laboratory parameters (see **Figure 7B**): (i) degree of thrombocytopenia in severe cases (all three diseases); (ii) eosinopenia (in COVID-19 and MIS-C, but not KD) and (iii) impaired cardiac contractility (unique to MIS-C; but not KD); (iii) an integrated analysis of serum cytokines and transcriptomics revealed that the proinflammatory MIP1α, TNFα and IL1 pathways are significantly induced in MIS-C compared to KD. In light of these findings, a rational approach to MIS-C treatment would be to combine the FDA-approved drugs anakinra^42^ and infliximab^43–45^. In fact, during the preparation of this manuscript a new study has already shown favorable outcome in MIS-C with the use of Infliximab^59^. Furthermore, our findings are consistent with the recently released guidelines by the American College of Rheumatology for initial immunomodulatory treatment of MIS-C^60^; it is noteworthy that these guidelines were released while this work was under review.

Finally, our findings reveal a pattern of MIS-C-defining molecular features (*IL15/IL15RA*, MIP1α, *TNF*α and *IL1* pathways) and clinical and laboratory parameters (thrombocytopenia, eosinopenia, and reduced myocardial function). For example, MIP1α elevation shows strong correlations with clinical features of disease (low PLT, high IL15 and low AEC) in MIS-C, but not KD. This suggests two things—(i) that despite shared proximal proximal pathways of immunopathogenesis (i.e., *IL15/IL15RA*-centric cytokine storm), the immunopathogenesis of KD and MIS-C may diverge distally; and (ii) that *IL15/IL15RA*, eosinopenia and thrombocytopenia may be inter-related phenomena in the setting of infection and inflammation. Platelets, besides their role in hemostasis, they are known to participate in the interaction between pathogens and host defense^61–63^. Persistent thrombocytopenia carried higher mortality in sepsis^64, 65^, and in COVID-19^66, 67^. Our analysis revealed a direct and unusually strong correlation between thrombocytopenia and eosinopenia in MIS-C. Eosinophils, on the other hand, as reviewed elsewhere^68^, have important antiviral properties, attributed to their granular proteins (e.g., eosinophil-derived neurotoxin, cationic protein) that display antiviral activities against single-stranded RNA viruses. Eosinophils can also support viral clearance^69^. Eosinopenia, in the setting of acute infection, has been found to be a direct response to infectious stimuli^70^, TLR4 ligands and chemotactic factors^71^, and has been considered a reliable diagnostic marker of infection^72^ in critically ill patients and a predictor of mortality^72–74^. Of relevance to the pediatric syndrome MIS-C, eosinopenia is encountered in about a 1/3^rd^ of the pediatric COVID-19 subjects^75^. It is noteworthy that eosinopenia (defined as an eosinophil count < 15 cells/μL and an eosinophil percentage < 0.25%) is a known poor prognostic factor for admissions into the pediatric ICU (hazard ratio [HR]: 2.96; P = 0.008^76^). It is possible that the two related clinical/laboratory parameters (low PLT and AEC) may be useful indicators of disease severity and prognosis in MIS-C and may guide decision-making in therapy and level of care in the hospital setting.

The strength of our study lies in the concurrent analysis of KD and MIS-C samples, our access to relatively large and independent cohorts of patients (in the case of KD), our ability to include age-matched pediatric healthy controls and febrile controls (non-KD and non-MIS-C, both pre-pandemic), and that all samples were drawn prior to the initiation of treatment. In doing so, this study overcomes some of the limitations of prior studies^12, 14, 16^. Another strength is the use of a set of *ViP* signatures (that were validated in COVID-19)^22^ and a KD-diagnostic signature^38^ as the computational framework to compare the two syndromes. Last, but not the least, the multi-omics approach used here on samples obtained from the same patients allowed us to predict and validate the prominent and invariant upregulation of one cytokine pathway (i.e., *IL15*) at both transcript and protein level. Notable limitations of our study include a relatively small sample size of MIS-C subjects (n = 12), limited number of publicly available MIS-C datasets for independent validation, and our inability to access cardiac tissue from KD and MIS-C subjects. Future studies on emerging datasets will enable rigorous validation of the analysis presented here.

## METHODS

### Subjects and sample collection

#### Kawasaki Disease (KD), multisystem inflammatory syndrome in children (MIS-C), febrile control (FC) Subjects

All KD subjects met the American Heart Association (AHA) criteria^77^ for complete or incomplete KD and subjects in this study were enrolled before the SARS-CoV-2 pandemic. Coronary artery Z-scores were classified according to the AHA 2017 guidelines as follows: normal < 2.0; dilated, 2 ≤ Z < 2.5; aneurysm: 2.5 < Z < 10.0; and giant aneurysm, ≥10.0. All MIS-C patients met the case definition from the Centers for Disease Control and Prevention. Subjects were enrolled prospectively with collection of acute, pre-treatment samples. Demographic and clinical data including echocardiography data and laboratory values were prospectively collected and entered into an electronic database. Febrile control patients had fever of at least three days duration and at least one mucocutaneous feature of KD including rash, conjunctival injection, or mucosal erythema. All were enrolled prior to the onset of the pandemic. Details of how the final diagnosis for the febrile control patients was adjudicated are outlined in supplementary methods. The characteristics of patient cohorts that were part of this study are included in **Supplementary Information 1**. The study protocol was reviewed and approved by the institutional review board at UCSD (UCSD # 14020). Written informed consent from the parents or legal guardians and assent from patients were obtained as appropriate. For all the deidentified human subjects, information including age, gender, and previous history of the disease, was collected from the chart following HIPAA guidelines. The study design and the use of human study participants was conducted in accordance to the criteria set by the Declaration of Helsinki.

#### Collection of blood samples and RNA isolation

Whole blood was collected into PAXgene® tubes (PreAnalytiX) for RNA and into red top tubes for serum before the initiation of any treatment (illness day ≤10) for the KD, MIS-C and FC subjects (illness day <15 for some FC subjects) and at the clinic visit (day 17-25 of Illness for subacute and day 289-3240 of Illness for late convalescent) for the KD subjects. RNA was extracted following manufacturer’s instruction (PAXgene Blood miRNA Kit). Serum was separated immediately by centrifugation and stored at −80°C until use.

#### Tissue samples

We obtained formalin-fixed, paraffin-embedded tissues from a 4-year-old female who died nine months after KD onset due to thrombosis of giant aneurysms. Written consent was obtained from the parents. The tissue sampling protocol was reviewed and approved by the institutional review board at UCSD (UCSD# 180587).

### Computational Methods

#### ViP and severe (s)ViP Signatures

ViP (Viral Pandemic) signature is derived from a list of 166 genes using Boolean Analysis of large viral infection datasets (<u>GSE47963</u>; <u>GSE113211</u>). This 166-gene signature was conserved in all viral pandemics, including COVID-19, inspiring the nomenclatures ViP signature. A subset of 20-genes classified disease severity called severe-ViP signature. To compute the ViP signature, first the genes present in this list were normalized according to a modified Z-score approach centered around StepMiner threshold (formula = (expr -SThr)/3*stddev). The normalized expression values for every probeset for 166 genes were added together to create the final ViP signature. The severe ViP signature is computed similarly using 20 genes. The samples were ordered finally based on both the ViP and severe-ViP signature. A color-coded bar plot is combined with a violin plot to visualize the gene signature-based classification.

#### Data analysis

Several publicly available microarrays and RNASeq databases were downloaded from the National Center for Biotechnology Information (NCBI) Gene Expression Omnibus (GEO) website ^78–80^. Gene expression summarization was performed by normalizing Affymetrix platforms by RMA (Robust Multichip Average)^81, 82^ and RNASeq platforms by computing TPM (Transcripts Per Millions)^83, 84^ values whenever normalized data were not available in GEO. We used log2(TPM + 1) as the final gene expression value for analyses. GEO accession numbers are reported in figures, and text. KD/MIS-C RNASeq datasets were processed using salmon. Batch correction was performed using ComBat_seq R package.

*StepMiner* analysis, Boolean analysis, *BECC (Boolean Equivalent Correlated Clusters) Analysis and methodologies for creation of heatmaps and hierarchical agglomerative clustering are detailed in Supplementary Online Methods*.

#### Statistical Analyses

Gene signature is used to classify sample categories and the performance of the multiclass classification is measured by ROC-AUC (Receiver Operating Characteristics Area Under The Curve) values. A color-coded bar plot is combined with a density plot to visualize the gene signature-based classification. All statistical tests were performed using R version 3.2.3 (2015-12-10). Standard t-tests were performed using python scipy.stats.ttest_ind package (version 0.19.0) with Welch’s Two Sample t-test (unpaired, unequal variance (equal_var=False), and unequal sample size) parameters. Multiple hypothesis correction were performed by adjusting *p* values with statsmodels.stats.multitest.multipletests (fdr_bh: Benjamini/Hochberg principles). The results were independently validated with R statistical software (R version 3.6.1; 2019-07-05). Pathway analysis of gene lists were carried out via the Reactome database and algorithm^85^. Reactome identifies signaling and metabolic molecules and organizes their relations into biological pathways and processes. Kaplan-Meier analysis is performed using lifelines python package version 0.14.6. Violin, Swarm and Bubble plots are created using python seaborn package version 0.10.1. The source code for Boolean analysis framework is available at https://github.com/sahoo00/BoNE and https://github.com/sahoo00/Hegemon.

### Experimental Methods

#### RNA Sequencing

For polyA capture: Total RNA was assessed for quality using an Agilent Tapestation 4200, and samples with an RNA Integrity Number (RIN) greater than 8.0 were used to generate RNA sequencing libraries using the TruSeq Stranded mRNA Sample Prep Kit with TruSeq Unique Dual Indexes (Illumina, San Diego, CA). Samples were processed following manufacturer’s instructions, modifying RNA shear time to five minutes. Resulting libraries were multiplexed and sequenced with 100 basepair (bp) paired end reads (PE100) to a depth of approximately 50 million reads per sample on an Illumina NovaSeq 6000. Samples were demuxltiplexed using bcl2fastq v2.20 Conversion Software (Illumina, San Diego, CA). For ribosomal/globin depletion: Library preparation and sequencing of 30 million 75 or 100 bp paired end reads was conducted using the Illumina’s TruSeq RNA Sample Preparation Kit, ribosomal and globin RNA depletion was performed using the Illumina® Ribo-Zero Gold kit and HiSeq 4000 at The Wellcome Centre for Human Genetics.

#### Human serum cytokines measurement

Human serum cytokines measurement was performed using the V-PLEX Custom Human Biomarkers from MSD platform. Human serum samples fractionated from peripheral blood of KD and MIS-C patients (all samples collected prior to the initiation of treatments) were analyzed using customized standard multiplex plates as per the manufacturer’s instructions.

#### Immunohistochemistry

Formalin-fixed, paraffin-embedded heart tissue sections from COVID19 and KD patients were stained anti-human IL15 receptor A polyclonal antibody (11:200 dilution; proteintech®, Rosemont, IL, USA; catalog# 16744-1-AP) and anti-human IL15 monoclonal antibody (1:10 dilution; Santa Cruz Biotechnology, Inc., Dallas, TX, USA; catalog# sc-8437) after heat-induced antigen retrieval with Tris buffer containing EDTA (pH 9.0). Sections were then incubated with respective HRP-conjugated secondary antibodies followed by DAB and hematoxylin counterstain (Sigma-Aldrich Inc., MO, USA; catalog# MHS1), and visualizing by Leica DM1000 LED (Leica Microsystems, Germany).

## DATA AVAILABILITY

Source data are provided with this paper. All data is available in the main text or the supplementary materials. The GEO datasets will be embargoed for one year after publication and released to readers upon request to the corresponding authors. Publicly available datasets used: <u>GSE109351</u>; <u>GSE73464</u>; <u>GSE15297</u>; <u>GSE68004</u>; <u>GSE18606</u>; <u>GSE9863</u>; <u>GSE63881</u>; <u>GSE73577</u>; <u>GSE16797</u>; <u>GSE166489</u>; <u>GSE126124</u>; <u>GSE166489</u>; <u>GSE167028</u>; <u>GSE11545</u>; <u>GSE116946</u>; <u>GSE100150</u>; <u>GSE147608</u>; <u>GSE122552</u>; <u>GSE79970</u>; <u>GSE149050</u>; <u>GSE153781</u>; <u>GSE148810</u>; <u>GSE75023</u>; <u>GSE27864</u>; <u>GSE21835</u>; <u>GSE57253</u>.

## CODE AVAILABILITY

The software codes are publicly available at the following links: https://github.com/sahoo00/BoNE ^86^ and https://github.com/sahoo00/Hegemon ^87^.

## ACKNOWLEDGEMENTS

This work was supported by the National Institutes for Health (NIH) grants R01-GM138385 (to DS), R01-AI141630 (to P.G), R01DK107585 (to S.D) and R01-AI155696 (to P.G, D.S and S.D), UCOP-RGPO (R01RG3780, R00RG2628 & R00RG2642 to P.G, D.S and S.D), iDASH U54HL108460, R01HL140898 (to JCB and AHT), and a UC San Diego Stem Cell Center Pilot award (to P.G, D.S and S.D). G.D.K was supported through The American Association of Immunologists Intersect Fellowship Program for Computational Scientists and Immunologists. Clinical sample collection was supported by the Patient Outcomes Research Institute (PCORI) CER-1602-3447 (to JCB), a Gordon and Marilyn Macklin Foundation grant (JCB), and a UC San Diego Stem Cell Center Pilot award (to P.G, D.S and S.D). This publication includes data generated at the UC San Diego IGM Genomics Center Utilizing an Illumina NOVASeq 6000 that was purchased with funding from a National Institutes of Health SIG grant (#S10 OD026929).

## AUTHOR CONTRIBUTIONS

JCB, ATH recruited the KD patients in cohort 1 and conducted the RNA seq studies; DS, JCB and PG conceptualized the project; CS, ATH, JoKa, JB, JCB, Pediatric Emergency Medicine Kawasaki Disease Research Group* recruited the KD and MIS-C patients in cohorts 2 and 3 and collected biological samples used in this study; GDK and CS carried out the serum cytokine analysis under the supervision of SD; SK and GDK carried out the immunohistochemical studies under the supervision of DS and PG; DS carried out all computational modeling and analyses and contributed all software used in this work; JiKi was responsible for the management of transcriptomic datasets and uploading to the NCBI. DS, GDK and PG prepared figures for data visualization; PG, GDK, C.S wrote the original draft of the manuscript; DS, GDK, AH, JoKa, CS, JCB edited and revised the manuscript. All co-authors approved the final version of the manuscript. DS, SD, JCB and PG supervised various parts of the project and secured funding; DS and PG administered the project.

## Captions of Supplementary Figures and Datasets

**Supplementary Information 1: Characteristics of patients in various cohorts (#1-4) used in this study**

**Supplementary Information 2: Excel sheet showing raw intensity values and the calculated concentration of individual cytokines, as measured using MesoScale Discovery assays.**

**Supplementary fig. S1** (related to Figure 2). **The KD-specific (13 gene) and the sViP (20 gene) signatures are independent.**

**Supplementary fig. S2** (related to Figure 4A). **Serum cytokine profiles in acute and convalescent Kawasaki Disease (KD).**

**Supplementary fig. S3** (related to Figure 5). **Correlation tests of cytokine levels, as determined by mesoscale and clinical/laboratory findings.**

