## Supplementary Online Materials for "An AI-guided signature reveals the nature of the shared proximal pathways of host immune response in MIS-C and Kawasaki disease"

**Conflict of interest statement:** Authors have declared that no conflict of interest exists.

¶ Equal contribution.
†Consortia: Pediatric Emergency Medicine Kawasaki Disease Research Group (see Supplemental Online Material)

*__Correspondence to__:

**Debashis Sahoo, Ph.D;** Assistant Professor, Department of Pediatrics, University of California San Diego; 9500 Gilman Drive, MC 0703, Leichtag Building 132; La Jolla, CA 92093-0831.

**Jane C. Burns, M.D.;** Director, Kawasaki Disease Research Center, Department of Pediatrics, University of California San Diego; 9500 Gilman Dr. MC 0641, La Jolla, CA 92093-0641

**Pradipta Ghosh, M.D.;** Professor, Departments of Medicine, and Cell and Molecular Medicine, University of California San Diego; 9500 Gilman Drive (MC 0651), George E. Palade Bldg, Rm 232, 239; La Jolla, CA 92093. Phone: 858-822-7633: Fax: 858-822-7636:

### CATALOG OF SUPPLEMENTARY MATERIALS


1. *Consortia authors*
2. *Detailed Methods*
    - Subjects and Sample Collection
    - Computational Methods
    - Experimental Methods
3. *Supplementary Figures and Legends (S1-S3)*
4. *Tables (1, 2)- uploaded as separate supplementary items.*
5. *Supplementary References*

### CONSORTIA:

A list of members of the "*Pediatric Emergency Medicine Kawasaki Disease Research Group*":

**Pediatric Emergency Medicine Kawasaki Disease Research Group:**

| Name | E-mail | Nature of Membership |
|---|---|---|
| John Kanegaye, MD | | Authorship Responsibility |
| Naomi Abe, MD | | Non-author members |
| Lukas Austin-Page, MD | | |
| Amy Bryl, MD | | |
| J. Joelle Donofrio-Ödmann, DO | | |
| Atim Ekpenyong, MD | | |
| Michael Gardiner, MD | | |
| David J. Gutglass, MD, MBI | | |
| Margaret B. Nguyen, MD | | |
| Kristy Schwartz, MD, MPH | | |
| Stacey Ulrich, MD | | |
| Tatyana Vayngortin, MD | | |
| Elise Zimmerman, MD | | |

### Detailed Methods

#### Subjects and sample collection:

***Kawasaki Disease (KD), multisystem inflammatory syndrome in children (MIS-C), febrile control (FC) Subjects*:**

All KD subjects met the American Heart Association (AHA) criteria[1] for complete or incomplete KD and subjects in this study were enrolled before the SARS-CoV-2 pandemic. Demographic and clinical data including echocardiography data and laboratory values were prospectively collected and entered into an electronic database. Coronary artery dimensions were described as Z-scores (internal dimension of the right and left anterior descending coronary arteries expressed as SDs from the mean normalized for body surface area). Coronary artery Z-scores were classified according to the AHA 2017 guidelines as follows: normal <2.0; dilated, $2 \leq Z < 2.5$; aneurysm: $2.5 < Z < 10.0$; and giant aneurysm, ≥10.0.

All MIS-C patients met the case definition from the Centers for Disease Control and Prevention. Subjects were enrolled prospectively with collection of acute, pre-treatment samples. Demographic and clinical data including echocardiography data and laboratory values were prospectively collected and entered into an electronic database.

Febrile control patients had fever of at least three days duration and at least one mucocutaneous feature of KD including rash, conjunctival injection, or mucosal erythema. All were enrolled prior to the onset of the pandemic. The final diagnosis for the control patients was adjudicated by a pediatric infectious disease specialist (JCB) and by a pediatric emergency room physician (JK) at least two months after initial presentation when testing results and clinical outcome were known. The final diagnoses of the 30 FC were defined by PCR or viral culture and included the following infections: 12 adenovirus, 5 EBV, 2 metapneumovirus, 3 rhinovirus, 3 influenza, 2 parainfluenza, 2 RSV, and 1 measles.

The study protocol was reviewed and approved by the institutional review board at UCSD (UCSD # 14020). Written informed consent from the parents or legal guardians and assent from patients were obtained as appropriate. For all the deidentified human subjects, information including age, gender, and previous history of the disease, was collected from the chart following HIPAA guidelines. The study design and the use of human study participants was conducted in accordance to the criteria set by the Declaration of Helsinki.

### Computational Methods:

#### *Data Availability*

Source data are provided with this paper. All data is available in the main text or the supplementary materials. RNA Seq datasets generated in this work have been deposited at NCBI GEO (GSE178491). The GEO datasets will be embargoed for one year after publication and released to readers upon request to the corresponding authors. Publicly available datasets used: GSE109351; GSE73464; GSE15297; GSE68004; GSE18606; GSE9863; GSE63881; GSE73577; GSE16797; GSE166489; GSE126124; GSE166489; GSE167028; GSE11545; GSE116946; GSE100150; GSE147608; GSE122552; GSE79970; GSE149050; GSE153781; GSE148810; GSE75023; GSE27864; GSE21835; GSE57253.

#### *ViP and severe (s)ViP Signatures*

ViP (Viral Pandemic) signature is derived from a list of 166 genes using Boolean Analysis of large viral infection datasets (GSE47963, GSE113211). This 166-gene signature was conserved in all viral pandemics, including COVID-19, inspiring the nomenclatures ViP signature [2]. A subset of 20-genes classified disease severity called severe-ViP signature [2]. To compute the ViP signature, first the genes present in this list were normalized according to a modified Z-score approach centered around StepMiner threshold (formula = (expr -SThr)/3*stddev). The normalized expression values for every probeset for 166 genes were added together to create the final ViP signature. The severe ViP signature is computed similarly using 20 genes. The samples were ordered finally based on both the ViP and severe-ViP signature. A color-coded bar plot is combined with a violin plot to visualize the gene signature-based classification.

#### *Data analysis*

Several publicly available microarrays and RNASeq databases were downloaded from the National Center for Biotechnology Information (NCBI) Gene Expression Omnibus (GEO) website

[3-5]. Gene expression summarization was performed by normalizing Affymetrix platforms by RMA (Robust Multichip Average)[6, 7] and RNASeq platforms by computing TPM (Transcripts Per Millions)[8, 9] values whenever normalized data were not available in GEO. We used log2(TPM +1) as the final gene expression value for analyses. GEO accession numbers are reported in figures, and text. KD/MIS-C RNASeq datasets were processed using salmon. Batch correction was performed using ComBat_seq R package.

#### *StepMiner Analysis*

StepMiner is a computational tool that identifies step-wise transitions in a time-series data.[10] StepMiner analysis is used to identify the threshold to convert continuous gene expression values into Boolean values (High/Low). StepMiner performs an adaptive regression scheme to identify the best possible step up or down based on sum-of-square errors. The steps are placed between time points at the sharpest change between low expression and high expression levels, which gives insight into the timing of the gene expression-switching event. To fit a step function, the algorithm evaluates all possible step positions, and for each position, it computes the average of the values on both sides of the step for the constant segments. An adaptive regression scheme is used that chooses the step positions that minimize the square error with the fitted data. Finally, a regression test statistic is computed as follows:

$$F\ stat = \frac{\sum_{i=1}^{n}(\widehat{X_i} - \bar{X})^2/(m-1)}{\sum_{i=1}^{n}(X_i - \widehat{X_i})^2/(n-m)}$$

Where $X_i$ for $i = 1$ to $n$ are the values, $\widehat{X_i}$ for $i = 1$ to $n$ are fitted values. m is the degrees of freedom used for the adaptive regression analysis. $\bar{X}$ is the average of all the values: $\bar{X} = \frac{1}{n} * \sum_{j=1}^{n} X_j$. For a step position at k, the fitted values $\widehat{X_i}$ are computed by using $\frac{1}{k} * \sum_{j=1}^{n} X_j$ for $i = 1$ to $k$ and $\frac{1}{(n-k)} * \sum_{j=k+1}^{n} X_j$ for $i = k + 1$ to $n$.

#### *Boolean Analysis*

**Boolean logic** is a simple mathematic relationship of two values, i.e., high/low, 1/0, or positive/negative. The Boolean analysis of gene expression data requires the conversion of expression levels into two possible values. The ***StepMiner*** algorithm is reused to perform Boolean analysis of gene expression data.[11] **The Boolean analysis** is a statistical approach which creates binary logical inferences that explain the relationships between phenomena. Boolean analysis is performed to determine the relationship between the expression levels of pairs of genes. The ***StepMiner*** algorithm is applied to gene expression levels to convert them into Boolean values (high and low). In this algorithm, first the expression values are sorted from low to high and a rising step function is fitted to the series to identify the threshold. Middle of the step is used as the StepMiner threshold. This threshold is used to convert gene expression values into Boolean values. A noise margin of 2-fold change is applied around the threshold to determine intermediate values, and these values are ignored during Boolean analysis.

**BECC (Boolean Equivalent Correlated Clusters) Analysis**

BECC analysis[12] is based on Boolean Equivalent [11] relationships, pair-wise correlation and linear regression analysis. BECC analysis identified ViP and severe-ViP signature using the BooleanNet statistic [2].

***Heatmaps and hierarchical agglomerative clustering***

Gene expression values were normalized according to a modified Z-score approach centered around *StepMiner* threshold (formula = (expr - SThr)/3*stddev). The samples were ordered according to average of the normalized gene expression values in the largest cluster along the Boolean path. The heatmap use red colors for the high values, white colors for the intermediate values and blue colors for low values. Gene names for few selected genes are highlighted on the left to show their expression patterns. Rows and columns are ordered based on hierarchical agglomerative clustering using python seaborn (version 0.10.1) clustermap function. Dendrograms are displayed for both rows and columns.

### **Supplementary Information (Tables):** *Separate uploads as excel sheets*

**Supplementary Information 1:** *Characteristics of patients in various cohorts (#1-4) used in this study*

**Supplementary Information 2:** *Excel sheet showing raw intensity values and the calculated concentration of individual cytokines, as measured using MesoScale Discovery assays.*

### Supplementary Figures

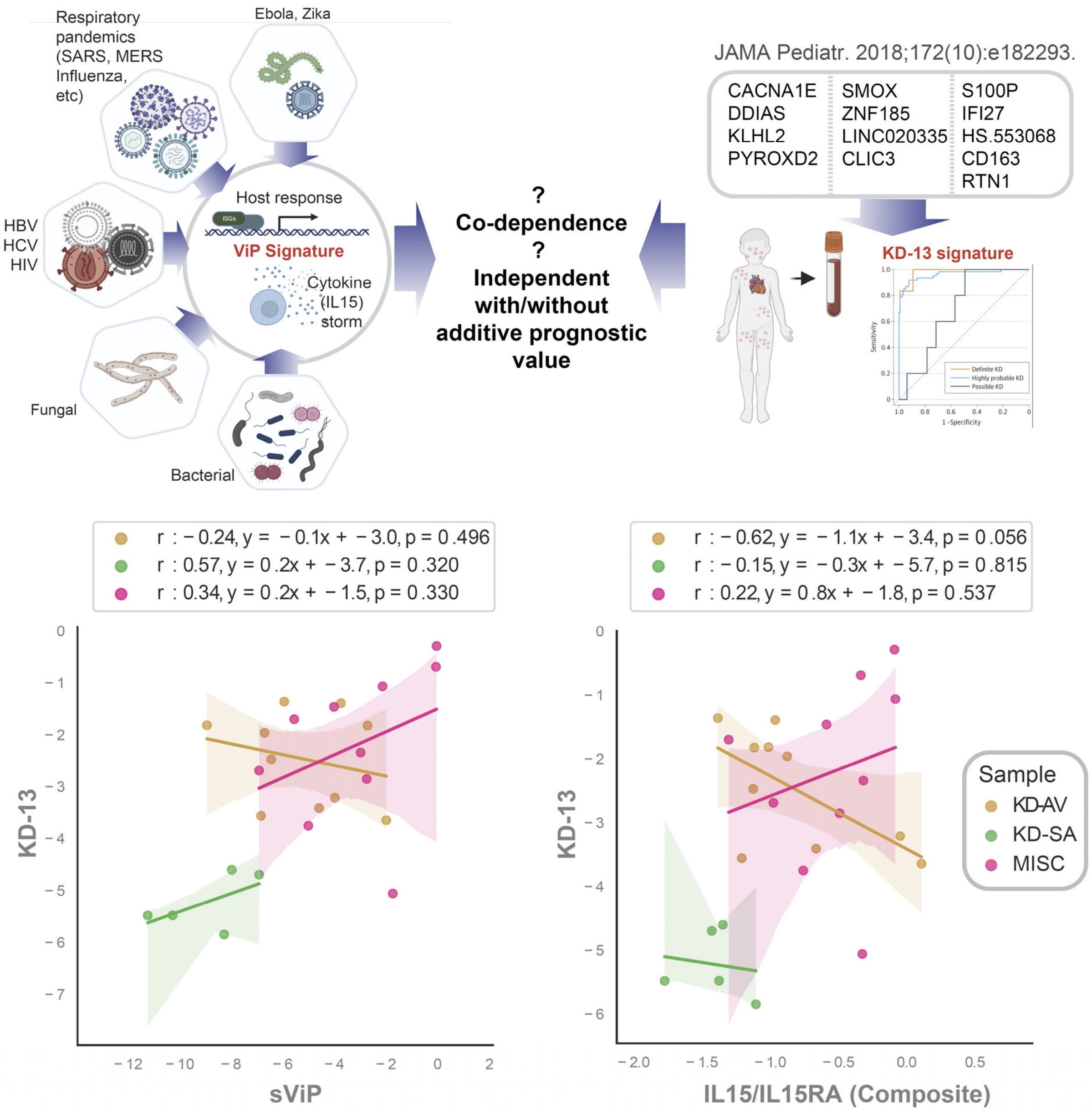


**Supplementary Figure S1** (related to Figure 2). **The KD-specific (13 gene)[13] and the sViP (20 gene) signatures are independent.** *Top*: Both sViP and KD-13 signatures are induced in the whole blood samples from MIS-C and KD subjects (**Figure 2C, H**), and here we asked if their induction may be co-dependent or independent using a correlation coefficient analysis. *Bottom*: Correlation tests between KD-13 signature (Y axis) and sViP (left, x axis) or IL15/IL15RA compositive score (right, x axis) were calculated and displayed as scatter plots using python seaborn lmplots. The confidence interval around the regression line is indicated with shades.

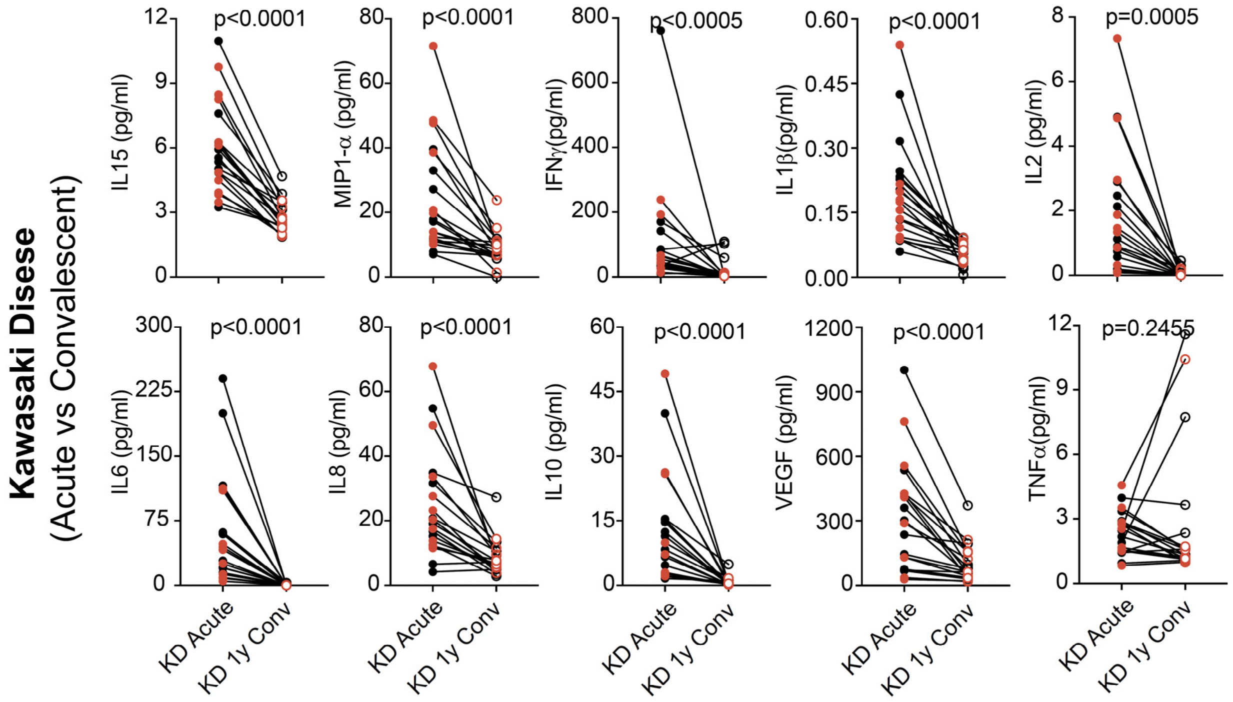


**Supplementary Figure S2** (related to Figure 4A). **Serum cytokine profiles in Kawasaki Disease (KD) in acute and convalescent disease.** Line plots show the changes in the levels of cytokines in paired samples from acute and convalescent visits of KD subjects. Wilcoxon matched pairs signed rank test was used for two parameter statistical analysis to test significance. Source data are provided as a Source Data file.

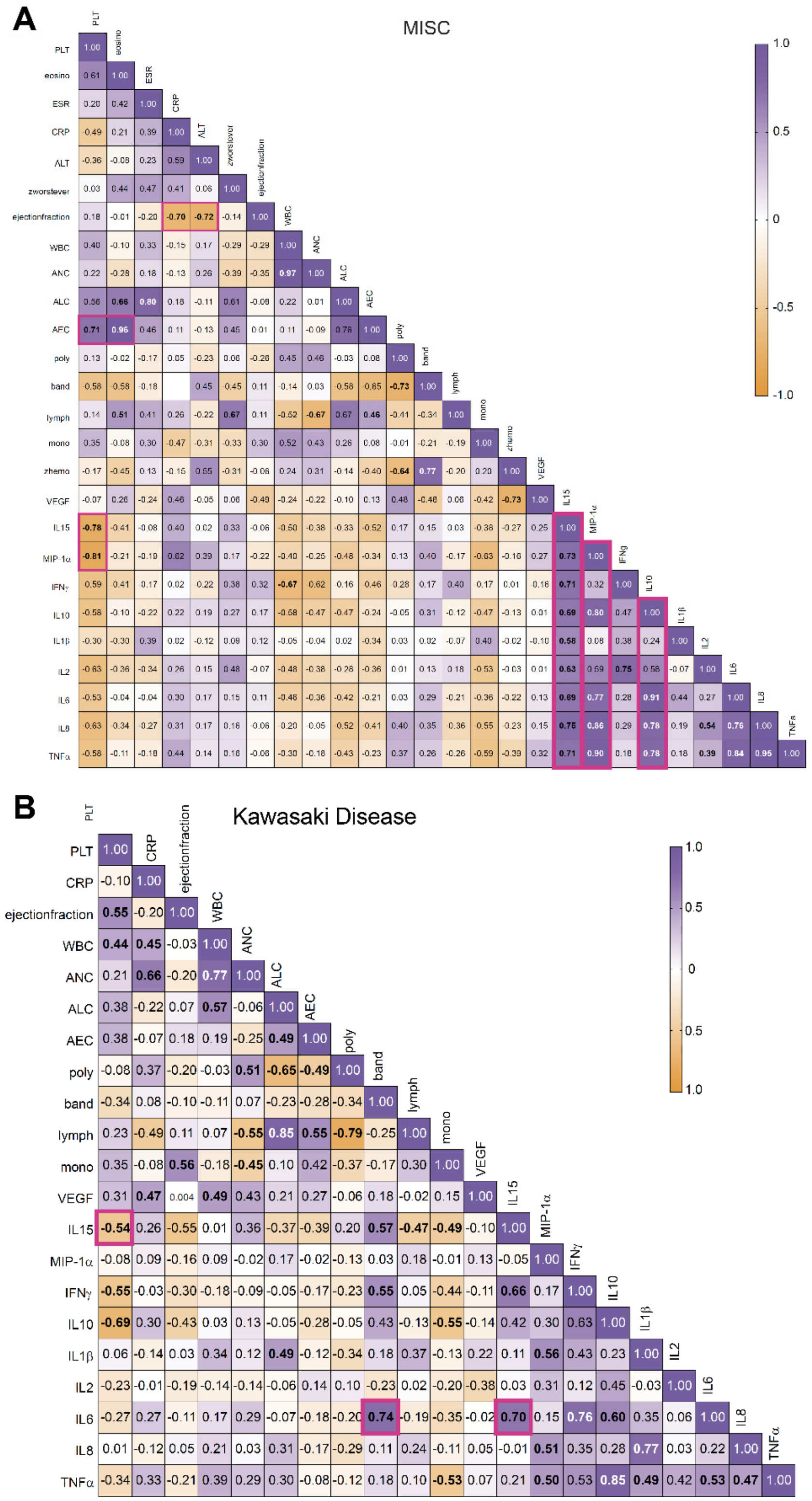


**Supplementary Figure S3** (related to Figure 5). **Correlation tests of cytokine levels, as determined by mesoscale and clinical/laboratory findings.** Heatmaps of correlation matrix are displayed for MISC (A) and KD (B). Correlation matrix and significance was determined using GraphPad Prism 9. Significant correlations, defined as those in which p values are < 0.05) are highlighted in bold fonts. Source data are provided as a Source Data file.
